# OMICmAge: An integrative multi-omics approach to quantify biological age with electronic medical records

**DOI:** 10.1101/2023.10.16.562114

**Authors:** Qingwen Chen, Varun B. Dwaraka, Natàlia Carreras-Gallo, Kevin Mendez, Yulu Chen, Sofina Begum, Priyadarshini Kachroo, Nicole Prince, Hannah Went, Tavis Mendez, Aaron Lin, Logan Turner, Mahdi Moqri, Su H. Chu, Rachel S. Kelly, Scott T. Weiss, Nicholas J.W Rattray, Vadim N. Gladyshev, Elizabeth Karlson, Craig Wheelock, Ewy A. Mathé, Amber Dahlin, Michae J. McGeachie, Ryan Smith, Jessica A. Lasky-Su

## Abstract

Biological aging is a multifactorial process involving complex interactions of cellular and biochemical processes that is reflected in omic profiles. Using common clinical laboratory measures in ~30,000 individuals from the MGB-Biobank, we developed a robust, predictive biological aging phenotype, *EMRAge*, that balances clinical biomarkers with overall mortality risk and can be broadly recapitulated across EMRs. We then applied elastic-net regression to model *EMRAge* with DNA-methylation (DNAm) and multiple omics, generating *DNAmEMRAge* and *OMICmAge,* respectively. Both biomarkers demonstrated strong associations with chronic diseases and mortality that outperform current biomarkers across our discovery (MGB-ABC, n=3,451) and validation (TruDiagnostic, n=12,666) cohorts. Through the use of epigenetic biomarker proxies, *OMICmAge* has the unique advantage of expanding the predictive search space to include epigenomic, proteomic, metabolomic, and clinical data while distilling this in a measure with DNAm alone, providing opportunities to identify clinically-relevant interconnections central to the aging process.

## MAIN

A major goal of aging research is to define biomarkers of aging that capture inter-individual differences in functional decline, chronic disease development, and mortality not identified through chronological age alone^1^. Both molecular and clinical data have been used to quantify various attributes of the biological aging process. Multiple molecular biomarkers of aging, or “clocks’’, have been developed as proxies for these hallmarks of aging^2^. These biomarkers have been variously based on telomere length^3^, neuro-imaging data^4–7^, immune cell counts^8^, and large-scale omics including DNA methylation (DNAm)^2,9–11^, metabolomics^12^, glycomics^13^, and proteomics^14–16^.

Over the last two decades, electronic medical records (EMR) have been widely used in clinical research, in particular for precision medicine, enabling deep phenotype mining from dense, comprehensive time-dependent data^17^. By utilizing comprehensive EMR data, we can capture clinical physiological changes over time that robustly illustrate phenotypic changes in real-time health status. Capitalizing on EMRs provides a unique opportunity to quantify the aging process in a reproducible way across clinical settings. While healthy aging encompasses both quality of life and life span, metrics of biological age have traditionally focused on using either clinical data to quantify quality of life^18,19^, or mortality risk to quantify life span^20^, resulting in biological phenotypes that are optimized to one of these attributes, while not fully capturing the other. With the wealth of data captured via EMRs, biological aging phenotypes that incorporate both dense clinical data and mortality can be created to synthesize these important attributes of aging into a single measure.

While clinical data are essential in creating aging phenotypes, connecting these phenotypes to the molecular underpinnings of the aging process is equally important. We do so by combining EMR data with comprehensive ‘omic profiling to assess the biological processes that ultimately govern aging. The strong molecular link between DNA methylation (DNAm) and the aging process has resulted in the widespread development and success of DNAm clocks with various biologic aging phenotypes aimed at reflecting clinical biomarkers (e.g. PhenoAge^18^), mortality (e.g. GrimAge^20^), and the rate of aging (e.g. DunedinPACE^21^).

Despite their broad use across both research and commercial settings, DNAm clocks have notable limitations. One such limitation is the difficulty of accurately reducing dimensionality due to the technical noise in measuring individual CpGs, which subsequently affects the precision ^22^. The heterogeneity of immune cell subsets is also a confounder of DNAm aging estimates, and whilst cellular deconvolution methods have been applied, to date immune deconvolution has considered a limited number of cell types^23,24^. Further, the inclusion of a CpG in a predictive aging clock does not necessarily imply causality nor functionality^25^, and identifying causal CpGs from DNAm clocks remains a challenging task, one that limits their biological potential.

Proteomics and metabolomics are more directly related to biological phenomena and may have utility as components of aging clocks. The proteome is altered by hallmarks of aging including loss of proteostasis, dysregulated nutrient sensing^26^, altered intercellular communication^27^ and cellular senescence^28^. Further, blood plasma contains circulating proteins derived from nearly all organs and cell types, making it possible to associate findings in peripheral blood with specific tissues and organs^21^. The metabolome not only provides critical information about metabolic processes, but it also provides measures of environmental exposures, including xenobiotics, that may be critically linked to the aging process. Further, the peripheral blood metabolome carries information from multiple tissues across the body, increasing the potential aging information of metabolomics compared to methylation and transcriptomic clocks of blood cells^29^.

Despite the important advantages of other omics, the development of transcriptomic, proteomic, and metabolomic clocks for biologic aging phenotypes has been limited. Initial work has demonstrated that while individual omics clocks share commonalities, each omic data type provides a distinct window of features that illustrate the aging process^30^, suggesting that the best and most clinically informative approach would harmonize combined information from multiple omic measurements to create an optimized aging clock. However, the integration of multiple omics into a multi-omic clock or to optimize and further inform upon the biological processes of improvement upon DNAm clocks remains an area of unfulfilled clinical potential.

Using ~30,000 participants from the Massachusetts General Brigham (MGB) Biobank, we developed and validated three distinct and clinically relevant measures of biological age: 1) *EMRAge*, a clinically-based and versatile death mortality predictor that can be broadly recapitulated across EMRs; 2) *DNAmEMRAge*, a DNA methylation aging biomarker trained to predict *EMRAge*; and 3) *OMICmAge*, the first DNA methylation-based multi-omic aging biomarker trained to predict *EMRAge*, that integrates proteomic, metabolomic, and clinical data through the use of Epigenetic Biomarker Proxies (EBP) (i.e., methylation surrogates) ^31–33^. By outperforming current methylation-based clocks in associations with chronic disease outcomes and mortality, we demonstrate the value of *DNAmEMRAge*, and *OMICmAge*, while further substantiating the biological relevance and value of integrating multiple omic data into one biological aging phenotype.

## RESULTS

### Overview of Study Design

To develop and validate *EMRAge, DNAmEMRAge*, and *OMICmAge,* we used participants in the Massachusetts General Brigham (MGB) Biobank. Individuals with available plasma and clinical data were used to develop *EMRAge* (n=30,884) (**Extended Figure S1**). A subset of these individuals who had available omic data were used to develop *DNAmEMRAge* and *OMICmAge* (MGB Aging Biobank Cohort (MGB-ABC), n=3,451). Finally, we validated these aging biomarkers using an independent cohort, the TruDiagnostic Biobank (n=12,666) (**Figure 1**). The additional clinical characteristics and demographics for these cohorts are described in **Extended Table S1**. Overall the population in the MGB Biobank cohort has a higher prevalence and broader range of comorbidities than individuals enrolled in the TruDiagnostic Biobank cohort.

**Figure 1.**
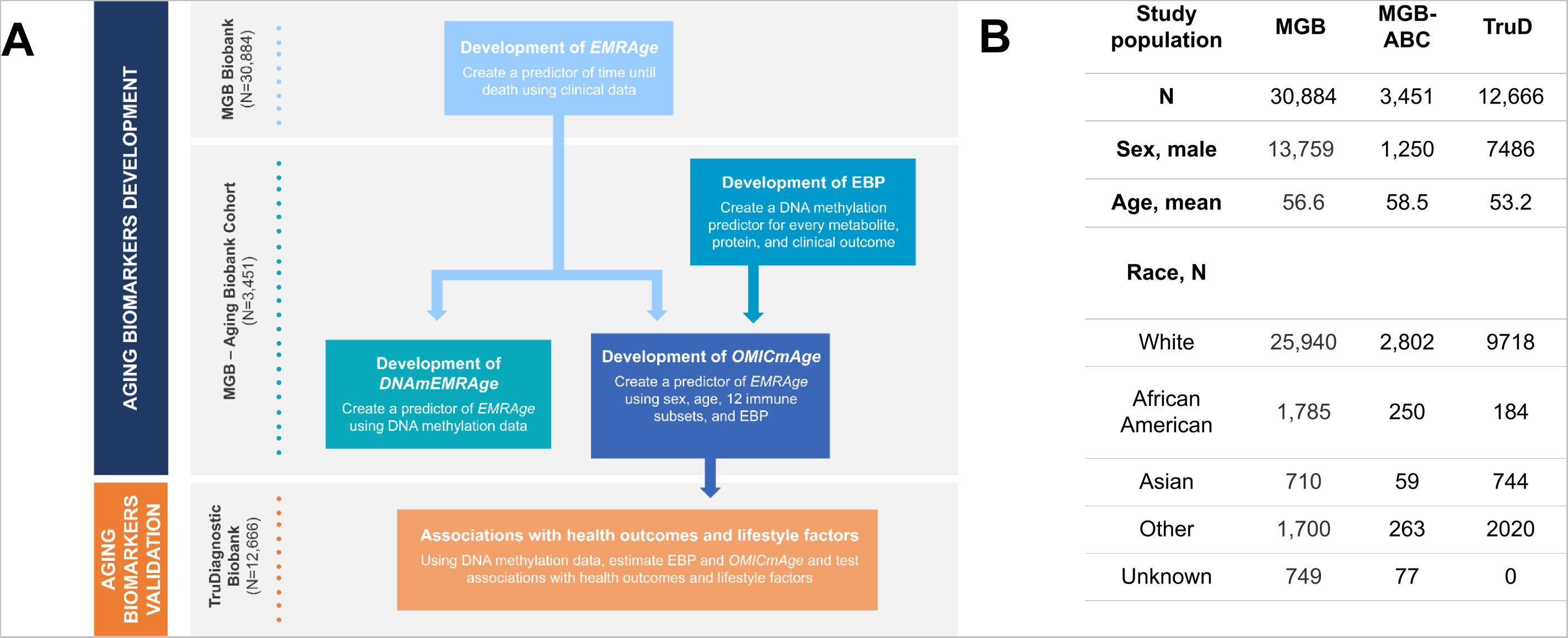
Overall study design. A) Workflow of the study. B) Description of the study population used in the study. MGB: Massachusetts General Brigham. MGB-ABC: MGB Aging Biobank Cohort. TruD: TruDiagnostic. EBP: Epigenetic Biomarker Proxy.

### Development of *EMRAge*

First, 30,884 individuals were apportioned randomly into training and testing sets using a 70:30 ratio. A Cox proportional hazards (Cox-PH) model was fitted in the training set to estimate the weightings of the 19 selected features (**Extended Table S7**). In a manner analogous to the GrimAge approach^20^, we converted the linear combination of estimated weights and predictor values into an “age” metric (**Method**). The Pearson correlation coefficient between *EMRAge* and chronological age was found to be greater than 0.75 (**Extended Figure S2**). We validated the *EMRAge* predictors by re-training the algorithm at four time points in 2-year increments: January 1st of 2008, 2010, 2012, and 2014. The four derived equations were then applied to participants (N=11,673) on January 1st of 2016. The Pearson correlations amongst these estimates were nearly 1, affirming the robustness of the *EMRAge* predictors (**Figure 2A**). **Figure 2B** shows that all aging-related health outcomes, including all-cause mortality, stroke, type-2 diabetes, chronic obstructive pulmonary disease (COPD), depression, other cardiovascular diseases (CVD), and any type of cancer, are significantly positively associated with higher *EMRAge*. The highest hazard ratio was seen for all-cause mortality (HR = 1.10 with 95% CI [1.10, 1.11]), followed by stroke and COPD. For the odds ratio, type-2 diabetes (OR = 1.08 with 95% CI [1.08, 1.09]) showed the highest values, followed by CVD and COPD. The training and testing sets showed very similar results for all the health outcomes. As illustrated in **Figure 2C**, participants with older *EMRAge* and older PhenoAge exhibit a markedly higher mortality risk than their younger counterparts. However, *EMRAge* more effectively discriminates between populations with high versus low survival probabilities, particularly among the oldest demographic. Furthermore, after adjusting for covariates, *EMRAge* has higher HR for all aging-related outcomes, with the exception of Type 2 Diabetes, when compared with *PhenoAge* (**Figure 2D**).

**Figure 2.**
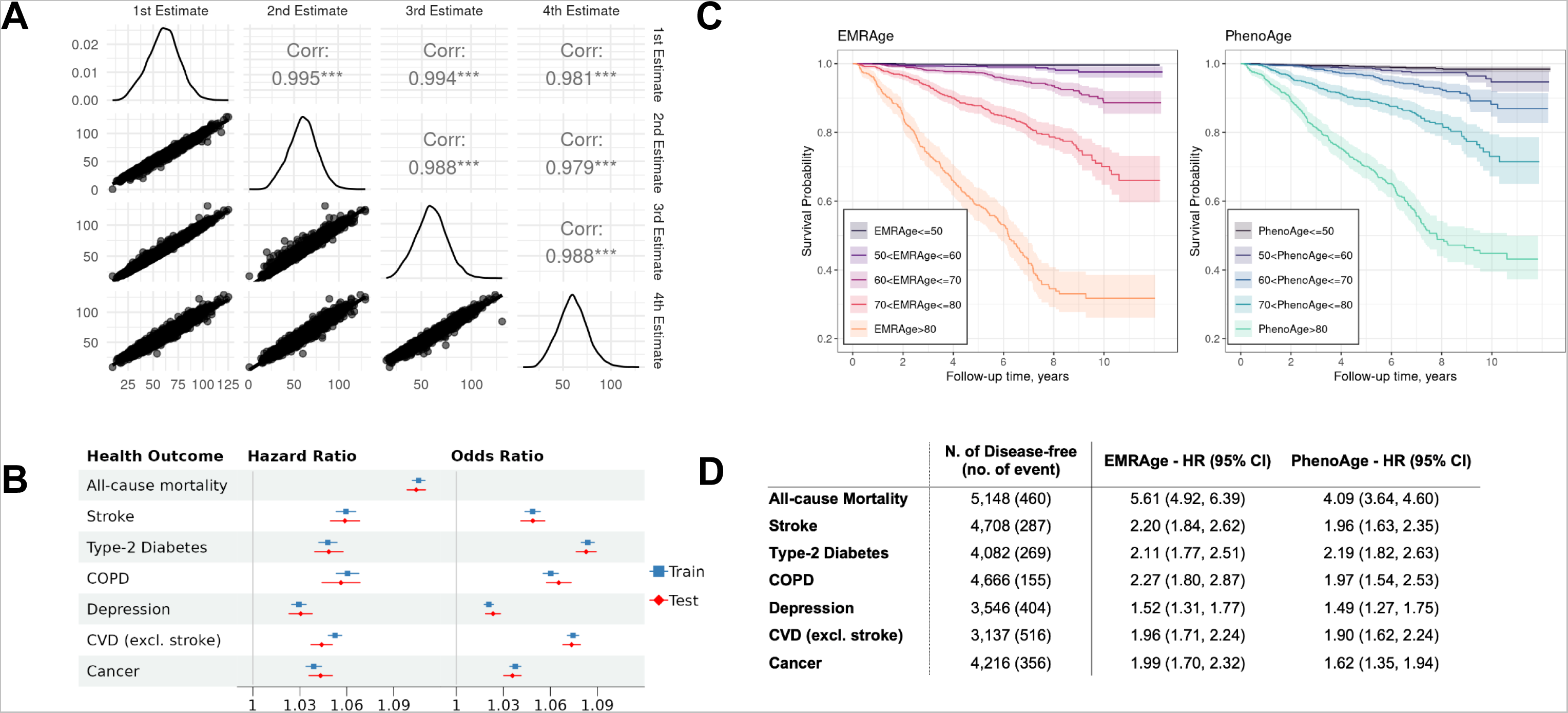
Development, Robustness, and Comparaters of *EMRAge*. A) Pairwise correlation between different 4 estimates of *EMRAge* at timepoints Jan 1st of 2008, 2010, 2012, and 2014. B) Forest plot of hazard ratio and odds ratio between *EMRAge* and aging-related health outcomes. C) Kaplan-Meier Plot of EMR Age vs. PhenoAge. D) Hazard ratios and confidence intervals of one standard deviation change to onset of aging-related diseases. These values were estimated in the testing dataset from the MGB Cohort (N = 5,148) adjusting for chronological age, sex, race, smoking status, and alcohol consumption.

### Development of *DNAmEMRAge*

After developing the *EMRAge* measure, we created a DNAm surrogate predictor of *EMRAge*, *DNAmEMRAge*, using DNA methylation data in an elastic net regression model (alpha=0.1) to select the CpGs that are most predictive of *EMRAge* (**Method**). The model for *DNAmEMRAge* included 1,097 CpG sites and age as predictors. A 25-fold cross validation showed an R^2^=0.827, suggesting good concordance in prediction. However, to further assess the agreement between *DNAmEMRAge* and the *EMRAge*, the data was resampled to identify a new training data set composed of samples used to generate the model and samples not in the model (N=2,762). Within the training data, the resulting *DNAmEMRAge* and *EMRAge* values showed high correlation (**Figure 3A**, N=2,762, R^2^=0.82, p<2.2e-16, Rho=0.91, p<2.2e-16). Finally, we also used a smaller test dataset which was not used for training to assess concordance; we find comparable correlations within this test set (N = 689, R^2^=0.83, p<2.2e-16, Rho=0.91, p<2.2e-16, see **Figure 3B**). Finally, the mean absolute error between *DNAmEMRAge* to *EMRAge* is 8.33 years in the training set and 8.50 years in the testing set and the intraclass correlation coefficient (ICC) was 0.995 (**Figure 3C**).

**Figure 3.**
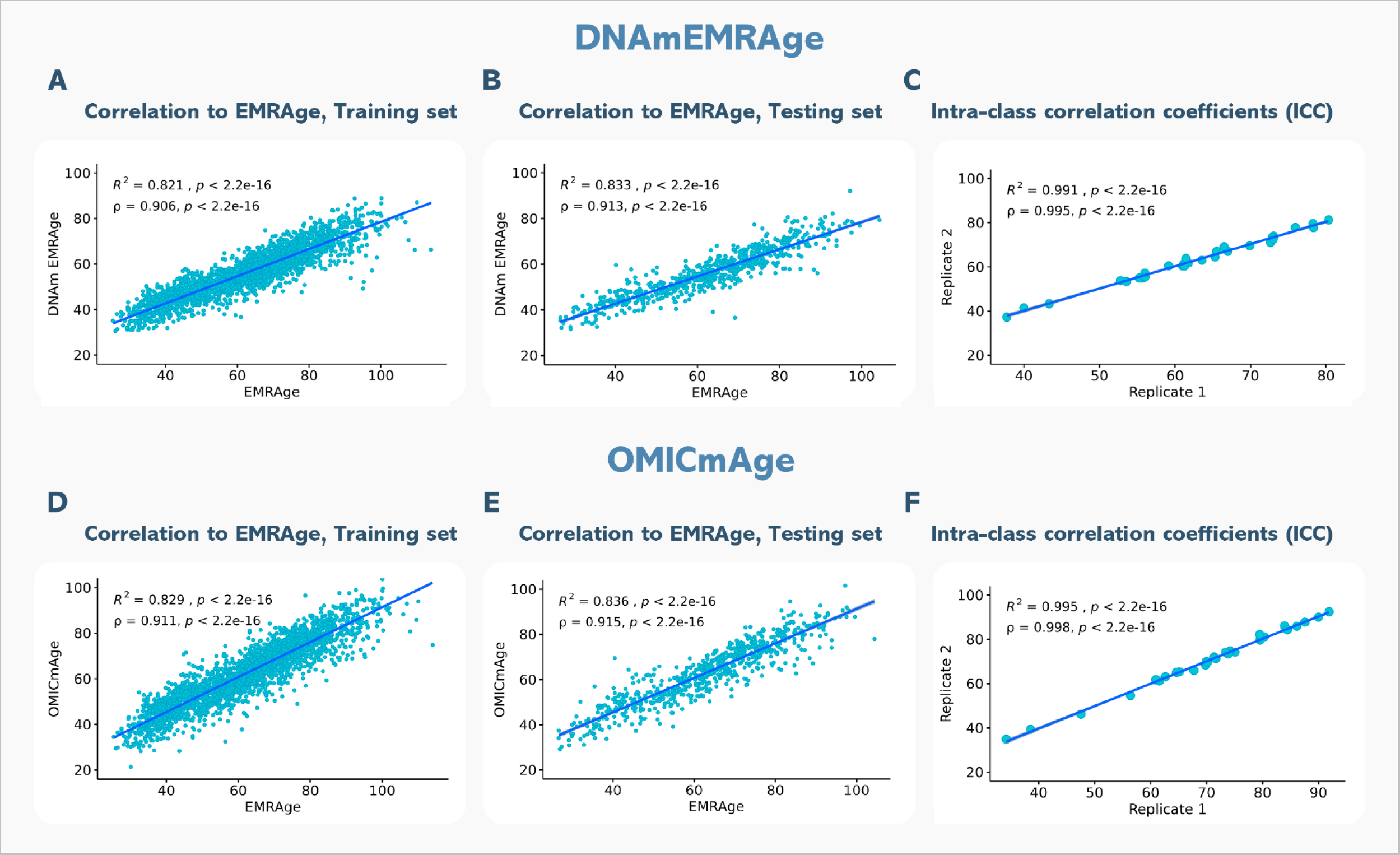
Correlation plots to *EMRAge* and intra-class correlation coefficients (ICC) for *DNAmEMRAge* and *OMICmAge*. A) Correlation between *DNAmEMRAge* and *EMRAge* in the training set (N=2762). B) Correlation between *DNAmEMRAge* and *EMRAge* in the testing set (N=689). C) Intra-class correlation coefficients for *DNAmEMRAge* using 30 replicates. D) Correlation between *OMICmAge* and *EMRAge* in the training set (N=2762). E) Correlation between *OMICmAge* and *EMRAge* in the testing set (N=689). F) Intra-class correlation coefficients for *OMICmAge* using 30 replicates.

### Development of *OMICmAge*

#### Metabolomic, Proteomic, and Clinical Epigenetic Biomarker Proxies (EBPs)

Untargeted global plasma metabolomic profiling was performed on the Metabolon platform. After preprocessing and scaling, the final dataset consists of 1,459 metabolites, that cover a broad range of metabolic pathways (**Extended Figure S3**), across 1,986 individuals, among which 1,691 were matched to methylation data. Global proteomic data were generated using the Seer SP100 platform, based on liquid chromatography mass spectrometry. The final processed dataset consisted of 2,805 non-unique and 536 unique protein groups (denoted as “*proteins”*) across 1,789 individuals, among which 1,475 were matched with methylation data. We further considered 46 clinical variables that have potential relationships with aging and aging-related outcomes. We selected proteins, metabolites, and clinical variables with a significant Pearson correlation (p<0.05) to *EMRAge* greater than 0.1, resulting in 299 metabolites, 110 proteins, and 25 clinical variables. We then generated Epigenetic Biomarker Proxies (EBPs) - epigenetic predictors for each selected metabolite, protein, and clinical variable - via an elastic net regression model. We retained all EBPs with a significant (p<0.05) Pearson correlation above 0.2 with their estimated metabolite/protein/clinical value. In total, 266 metabolite EBPs, 109 protein EBPs, and 21 clinical EBPs were retained, totaling 396 EBPs to be included as features in the predictive model for *OMICmAge* (**Extended Table S2**). *OMICmAge* was then generated by integrating proteomic, metabolomic, and clinical data into a DNA methylation clock.

#### Predictive model for the OMICmAge

*OMICmAge* was generated via a penalized elastic net regression model of *EMRAge* that included methylation CpG values, relative percentages of 12 immune cell subsets, 396 EBPs (**Extended Table S2**), age and sex as features in the model. This model retained 990 CpGs, 40 EBPs (16 protein EBPs, 14 metabolite EBPs, and 10 clinical EBPs) (**Figure 4A**) and age as significant predictors of *EMRAge* with varying weightings in the final model. Interestingly, the model did not retain any of the immune cell subsets after penalization. We tested an independent model including them as unpenalized features, but results did not change substantially. Thus, we continued with the model where all the features were penalized. **Figure 3** shows the correlation between *EMRAge* and *OMICmAge* in the training (N = 2,762, R^2^=0.83, p<2.2e-16; rho=0.91, p<2.2e-16) and testing sets (N = 689, R^2^=0.84, p<2.2e-16; rho=0.92, p<2.2e-16), as well as the ICC using 30 replicates (0.998). In terms of error, the mean absolute error between *OMICmAge* and *EMRAge* was 4.96 years in the training set and 4.97 years in the testing set, which was notably lower than the mean absolute error for *DNAmEMRAge* (8.33 and 8.50, respectively).

**Figure 4.**
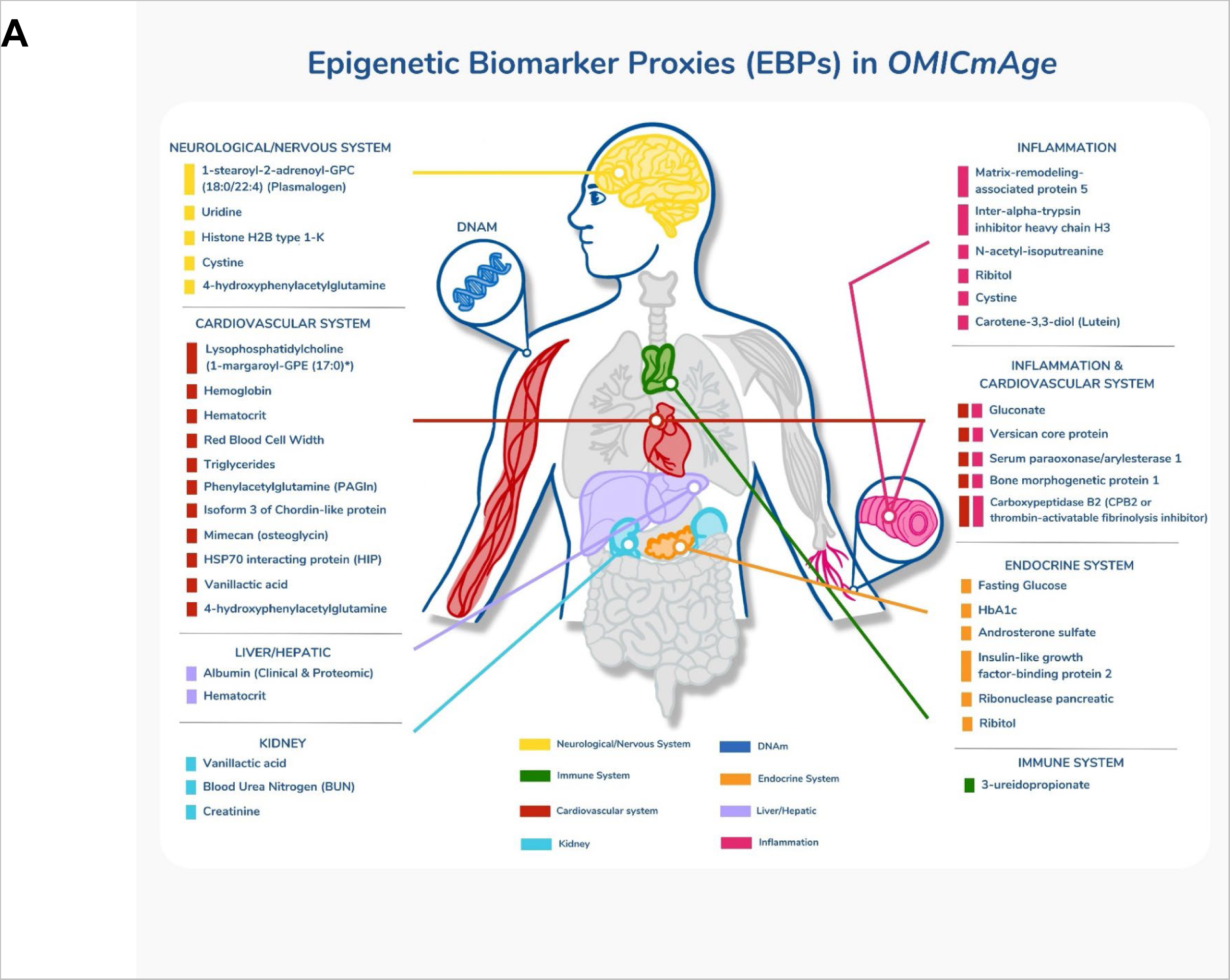
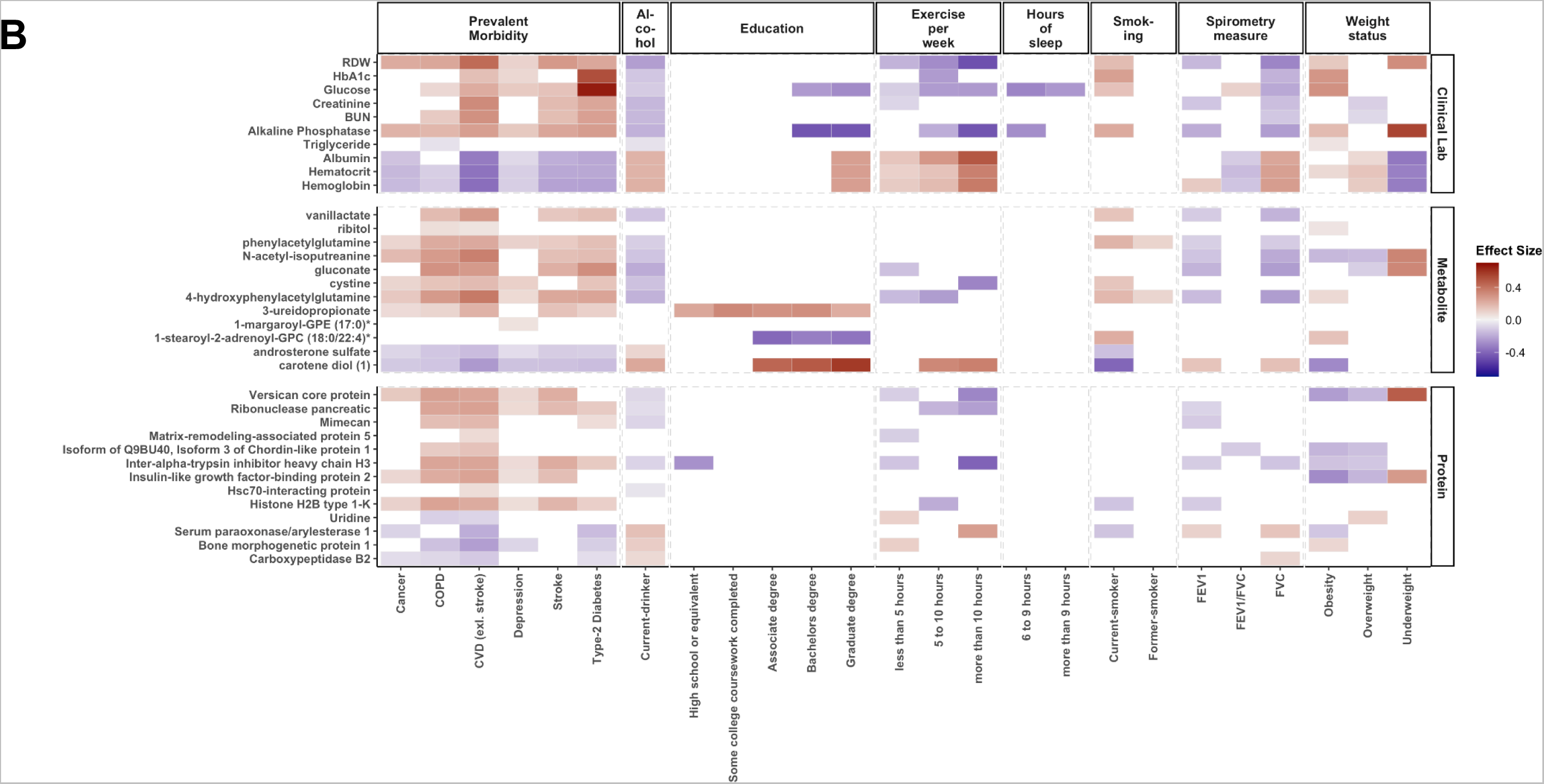
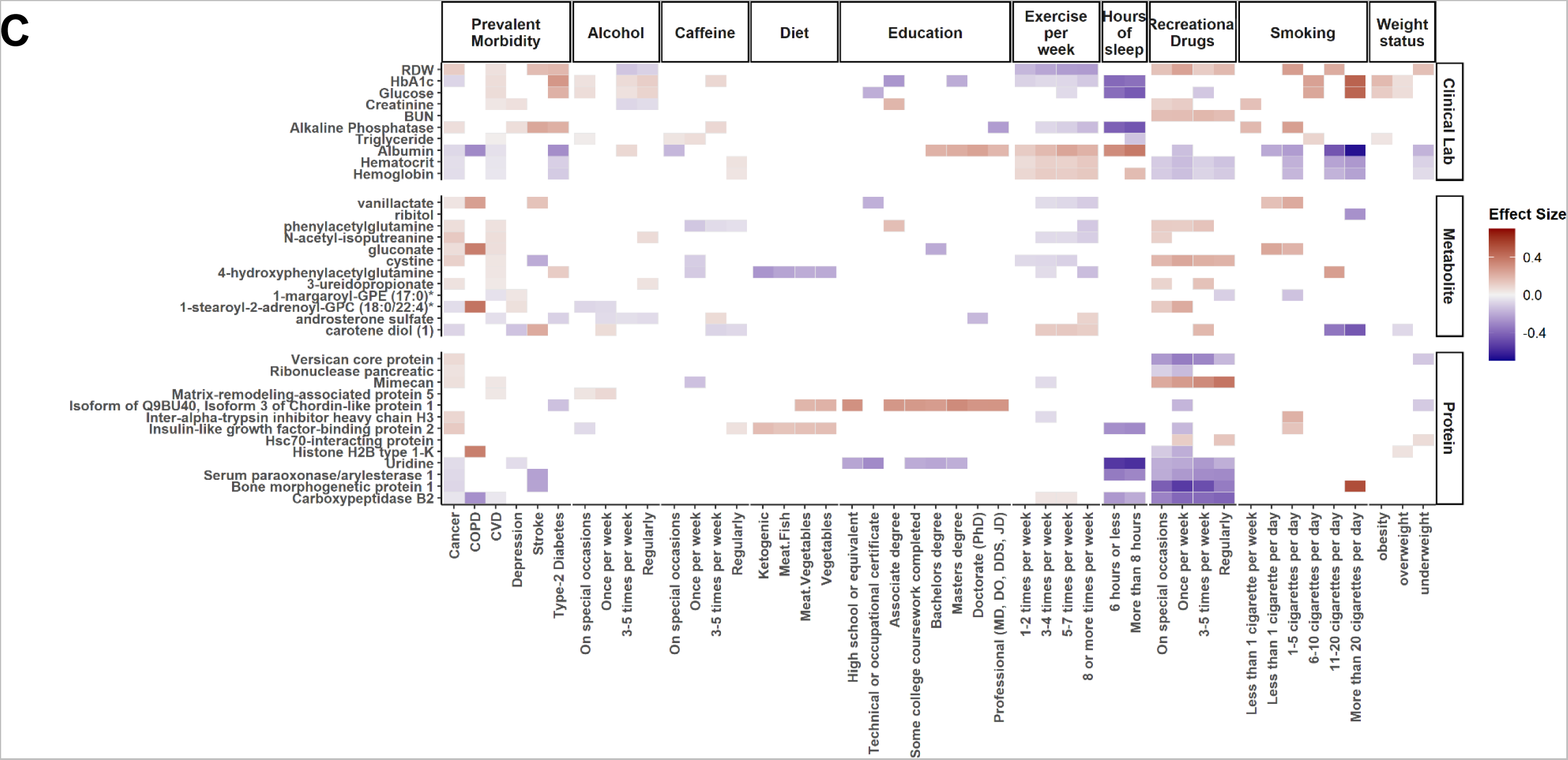
Epigenetic Biomarker Proxies (EBPs) included in the *OMICmAge*. A) EBPs selected in *OMICmAge* after penalization of the *OMICmAge* segmented by a system approach. Each color represents one of the 7 systems. B) Association of EBPs to diseases and lifestyle factors in the MGB-Aging Biobank Cohort. C) Association of EBPs to diseases and lifestyle factors in the TruDiagnostic biobank. We evaluated the association between each EBP and 6 major diseases (Depression, Stroke, COPD, Cancer, CVD, and Type-2 Diabetes Mellitus) and multiple lifestyle factors, including alcohol consumption, education level, and exercise per week, among others. The strength of the color is proportional to the estimate of the linear regression adjusted by age, sex, ethnicity, BMI, and tobacco smoking. In red, positive associations. In blue, negative associations.

#### Inferring the underlying biology from selected EBPs in the OMICmAge

A total of 40 EBPs spanning 8 biological systems are incorporated into *OMICmAge* and they predominantly represent cardiovascular, inflammatory, and endocrine systems; however, several of the selected metabolites and proteins EBPs do not have current clinical applications. We observed strong correlations between several of the selected EBPs and actual clinical values (e.g. rho=0.66 and 0.63 for EBP(CRP) and EBP(HbA1C) respectively). We also found significant associations between traditional disease biomarkers and EBPs, including positive associations between EBP(glucose), EBP(HbA1c) and type 2 diabetes and negative associations between EBP(FEV1) and COPD. Together, the directionality of these EBPs and their associated disease states are consistent with clinical disease biomarkers in both cohorts (**Figure 4B-C**). We observed more significant disease associations with EBPs in the MGB-ABC, which is likely due to the decreased health in MGB-ABC (**Extended Table S1**). We observed multiple associations between lifestyle factors and EBPs (**Figure 4B-C**). Overall, the direction of effect for these associations was consistent with the known biological knowledge for the relationship between the actual metabolite, protein, and clinical measurements and the specific chronic disease and lifestyle features being tested. This suggests that the EBPs are effective at capturing the underlying biological relationship of the original biomarkers.

#### Comparison of OMICmAge to previous epigenetic biomarkers of aging

We compared *DNAmEMRAge* and *OMICmAge* to previous epigenetic clocks in terms of their relationship with immune cell subsets, the CpGs sites included in the predictive model, the relationship with age-related disease outcomes, and five- and ten-year mortality. We generally observed consistent correlations between all epigenetic clocks and immune subsets; however, we observed stronger correlations with sex for both *OMICmAge* and *DNAmEMRAge* (R=0.28, p-value = 0.02, and R=0.36, p-value = 0.009, respectively) in comparison to previous clocks (**Extended Figure S4B**). There was minimal overlap between the CpG sites selected for estimating *DNAmEMRAge* and *OMICmAge* compared with previous clocks (**Figure 5A**); *DNAmEMRAge* and *OMICmAge* had 660 and 657 unique CpG sites respectively. Interestingly 411 CpG sites are shared between these two clocks. While PhenoAge and Horvath clock share 50 CpG sites and Horvath and Hannum share 29, the maximum number of probes shared between *OMICmAge* and any previous clock is 3. We also compared the prevalence and incidence of age-related disease outcomes between *DNAmEMRAge*, *OMICmAge,* and other aging clocks in the MGB-ABC and Tru Diagnostic Biobank cohorts (**Figure 5B, Extended Table S3**). For MGB-ABC *OMICmAge* or *DNAmEMRAge* had the highest ORs for type-2 diabetes, stroke, CVD and depression whereas *PCGrimAge* had the highest OR for COPD and the first-generation clocks (*PCHorvath pan tissue*, *PCHorvath* skin and blood, and *PCHannum*) had the highest ORs for cancer. Regarding HRs in MGB-ABC, *OMICmAge* or *DNAmEMRAge* had the highest HRs for type-2 diabetes, stroke, CVD, depression, COPD and all-cause mortality whereas *PCGrimAge* and *PCHorvath pan tissue* had the highest HRs with cancer. We observed comparable findings for prevalent disease associations in the Tru Diagnostic Biobank cohort, with *OMICmAge* or *DNAmEMRAge* having the highest ORs for all comorbidities except COPD where *PCGrimAge* had the highest observed OR. We also calculated the Area Under the Curve (AUC) for 5-year and 10-year survival using prediction classifiers for *OMICmAge, DNAmEMRAge, PCGrimAge*, and chronological age (**Figure 5C**). *DNAmEMRAge* showed the highest AUC values (5-year: 0.894, 10-year: 0.889), followed by *OMICmAge* with very similar values (5-year: 0.889, 10-year: 0.874). PC GrimAge and chronological age had AUC values that were approximately 5 percent less accurate than either *DNAmEMRAge* or *OMICmAge*.

**Figure 5.**
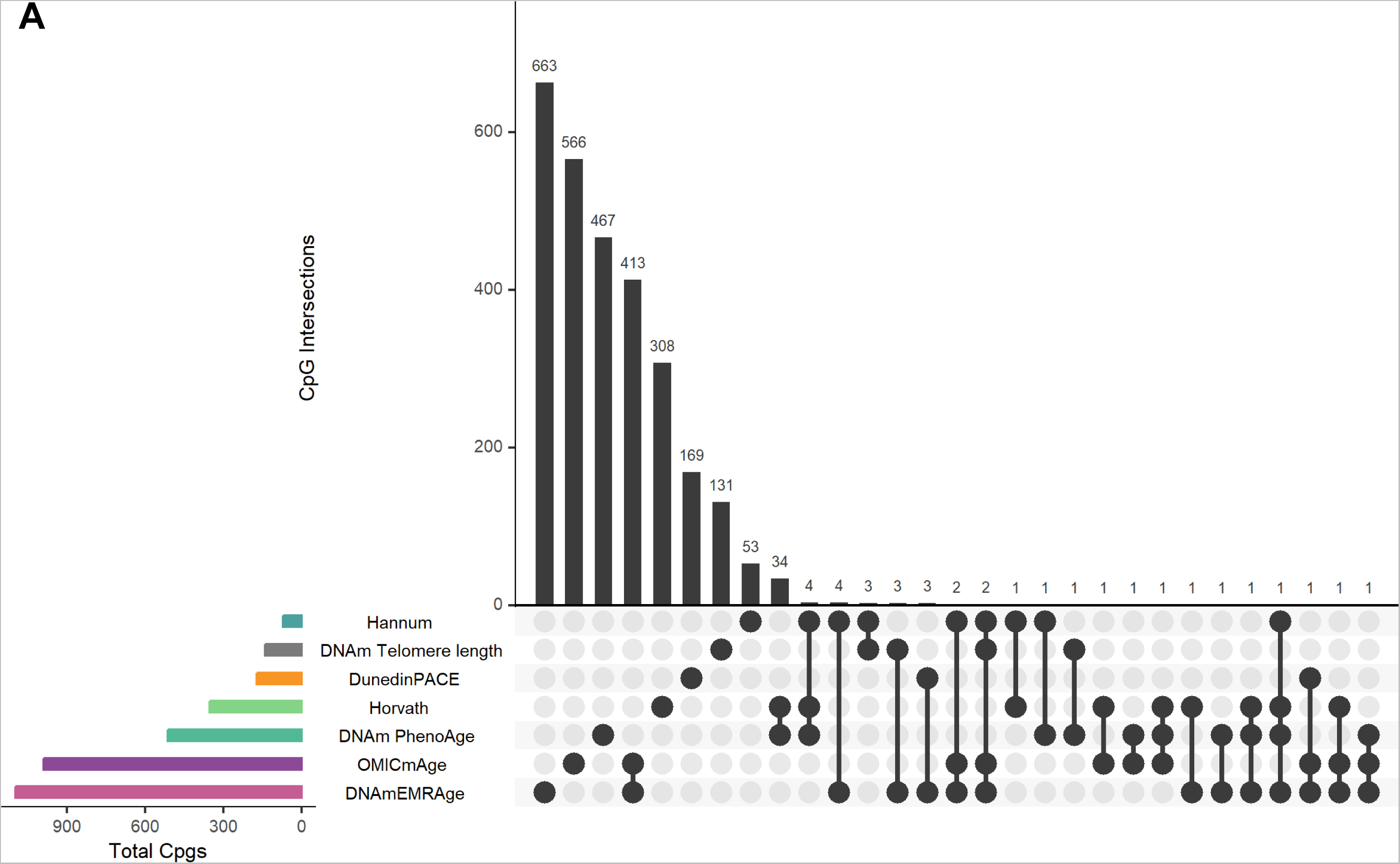
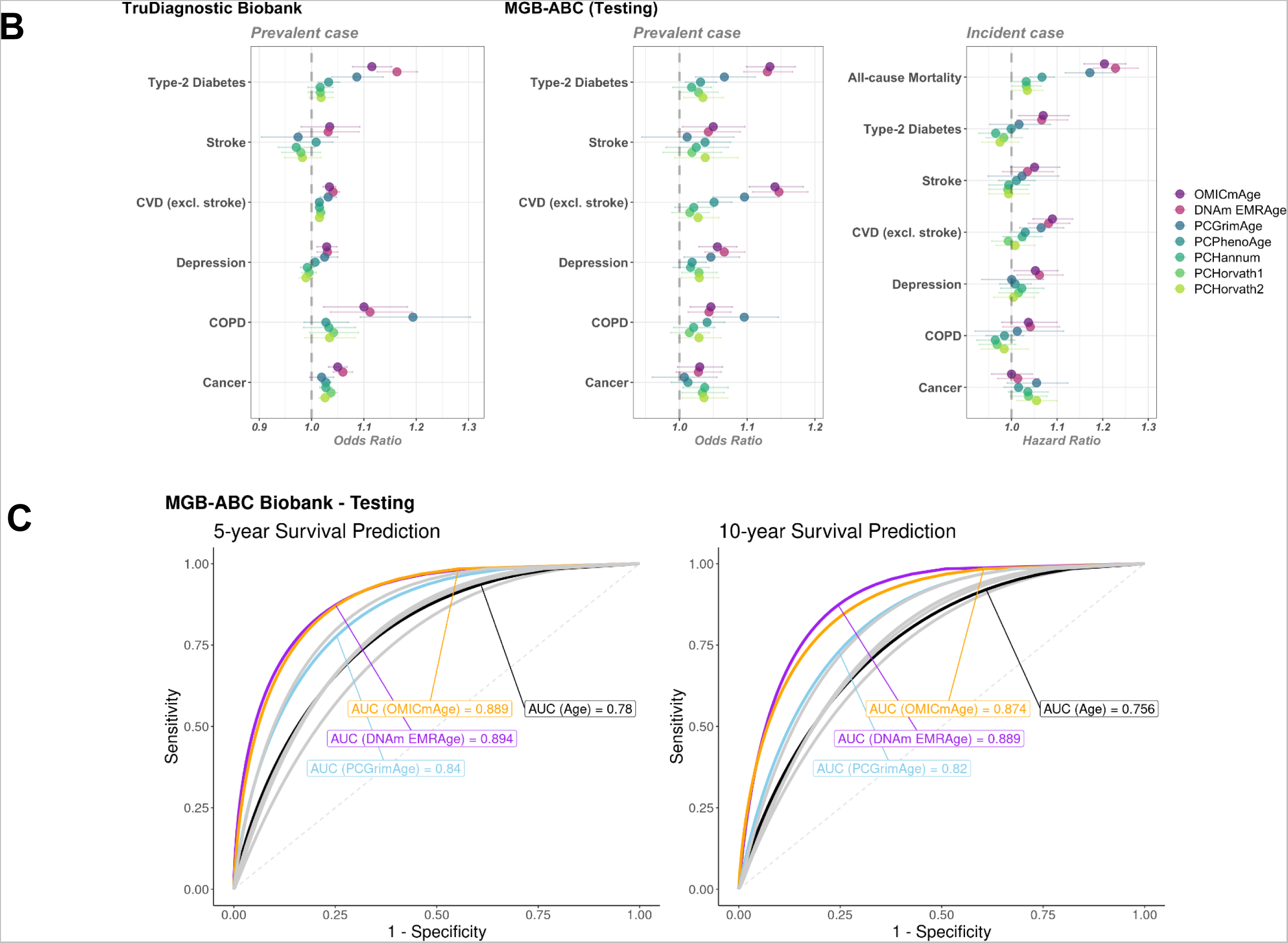
Comparison of *OMICmAge* and *DNAmEMRAge* to previously established aging biomarkers. A) Intersection of predictive CpG sites included in the previously published epigenetic clocks, DNAmEMRAge, and OMICmAge. The horizontal bars represent the total number of CpG sites included in each epigenetic clock. The vertical bars represent the number of unique or shared CpG sites between clocks. PhenoAge refers to the DNA methylation version. B) Horizontal errorbar plot of odds/hazard ratios of each methylation clock to aging-related diseases in TruDiagnostic Biobank cohort or testing set of MGB-ABC cohort. C) ROC curves for 5-year and 10-year survival prediction classifiers utilizing prior methylation clocks or chronological age. The orange line represents OMICmAge, the purple line represents DNAm EMRAge, the light blue line represents PCGrimAge, the remaining grey lines represent other PC aging clocks.

Using MGB-ABC, we observed strong positive associations between *OMICmAge* and male sex, tobacco smoking, chronological age, and body mass index (BMI) while we observed significant negative associations with physical activity and higher education (**Figure 6A**). In the TruDiagnostic cohort (**Figure 6B**), we observed similar estimated effects for sex, tobacco smoking, chronological age, BMI, and physical activity. When evaluating other lifestyle factors, recreational drug consumption was associated with a higher *OMICmAge* while high sexual activity, fish oil supplementation, and antioxidants were associated with a lower *OMICmAge* (**Extended Table S4**).

**Figure 6.**
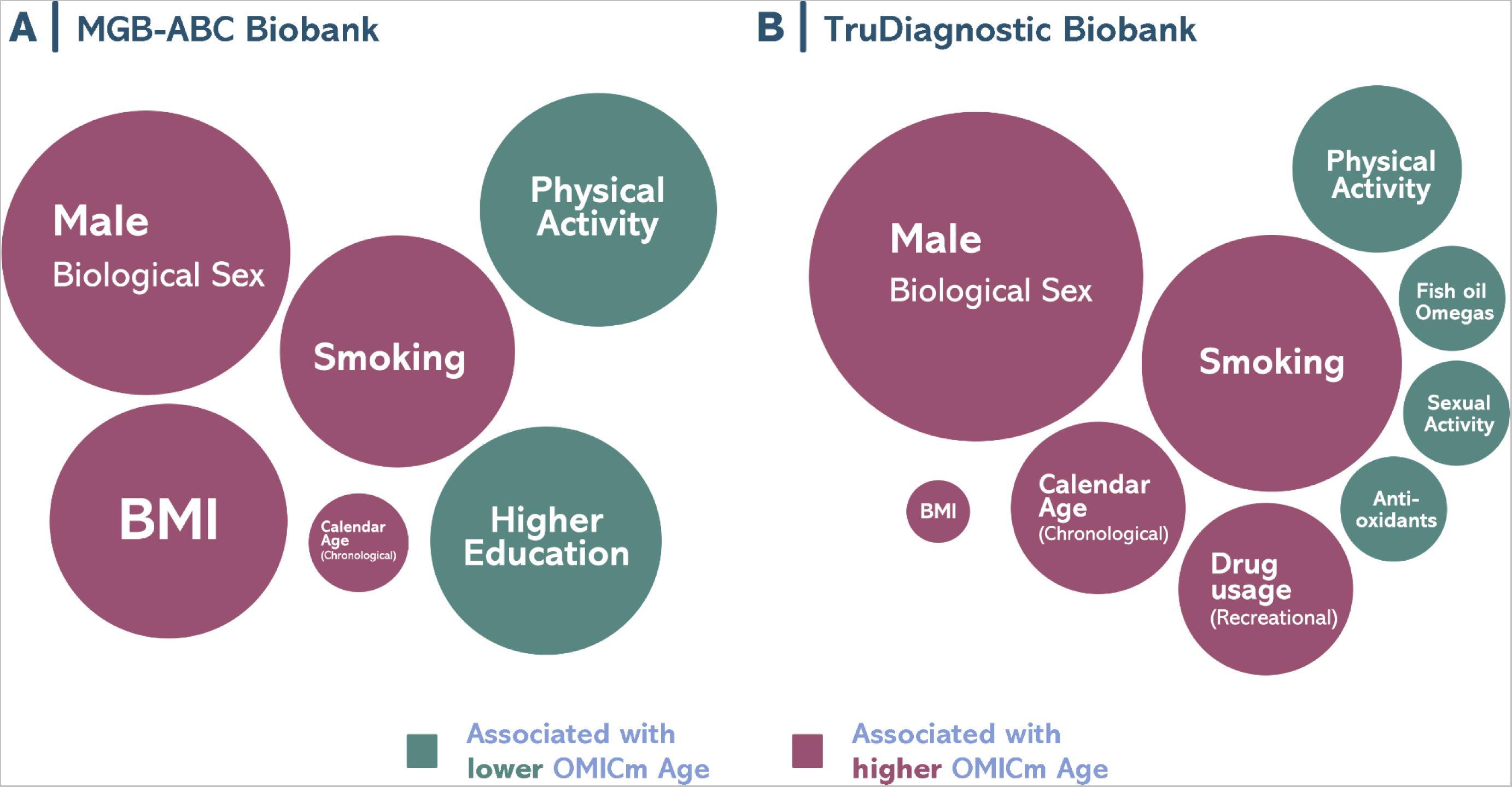
Bubble plot for the representation of the lifestyle factors associated with *OMICmAge*. A) MGB-ABC Biobank. B) TruDiagnostic Biobank. Visual representation of the effect sizes found with the OMICmAge. Circle diameter represents the calculated estimated value of the factor. Significant positive associations are represented in green whereas significant negative associations in red. All the associations are adjusted by chronological age, biological sex, ethnicity, body mass index (BMI), and tobacco use.

## DISCUSSION

Previously defined aging biomarkers have been developed using either clinical data or mortality prediction models alone^22,23^. Recognizing that both clinical measures that reflect overall health and the overall risk of death are critical yet distinct attributes of aging, we developed *EMRAge*, a hybrid aging phenotype that distills measures of both health and wellness and mortality into a single measure, as demonstrated by the robustness of *EMRAge*. We created a predictive model of time until death using 27 clinical EMR measures and mortality data from ~30,000 individuals in the MBG Biobank, spanning up to 30 years. When assessing mortality, we found that *EMRAge* demonstrates more accurate mortality risk prediction than either chronologic age or *PhenoAge* over a 10-year period. By creating multiple predictive models of *EMRAge* at different points in time in the EMR, we found strong reproducibility of *EMRAge* as an aging phenotype trained with common laboratory measures. We then created both DNAm and multi-omic biomarkers for aging for *EMRAge, DNAmEMRAge,* and *OMICmAge*. We demonstrate that both *DNAmEMRAge* and *OMICmAge* have strong associations with chronic diseases and mortality and outperformed current DNAm aging biomarkers. Importantly, we observe strong associations with multiple age-related diseases across two diverse cohorts representing poor to moderate overall health (MGB-ABC) and moderate to excellent overall health (TruDiagnostic Biobank) as demonstrated in **Extended Table S1**. Furthermore, we demonstrate marked improvements in the accuracy of both 5- and 10-year mortality risk. While both *DNAmEMRAge* and *OMICmAge* have similar overall performance, through the use of EBPs, the generation of *OMICmAge* has expanded the predictive search space to consider age-relevant proteins, metabolites, and clinical data while being generated through DNA methylation data alone. This expanse of biological input via multiple omics provides new translational opportunities for identifying clinically-relevant interconnections that are central to the aging process.

To date, aging clocks have been generated predominantly using singular omic data types, with varying degrees of accuracy^31,37^. It is clear that DNA methylation is one of the strongest predictors of aging phenotypes^26^; however, other omic data - in particular proteomics and metabolomics - have also demonstrated strong predictive accuracy and add the distinct advantage of providing more tangible biological insights into the aging process^24^. While the epigenome is a central mechanism for aging, the biological functions of epigenetic perturbations are often less clear to identify. In contrast, proteins capture a broad range of age-related biology, including immune function and inflammatory processes that are often well-understood and have clear clinical implications for treatment and/or modification. Changes in oxidative stress, hormones, and lipid profiles are just a few examples of the metabolic processes captured via the metabolome that reflect specific biology relevant for aging processes^27^. As such, by including the metabolome, proteome, epigenome, and clinical data into predictive space for *OMICmAge*, our modeling approach captures the aging processes on multiple levels of systems biology that may further elucidate relevant biological aging functions.

One ongoing challenge with multi-omic approaches is the complexity of integrating different omic data together and the subsequent interpretation of the findings. Moreover, final biological models that include multiple omic data types are most often impractical from a clinical perspective due to high costs, logistical difficulties, and time delays from multiple assay measurements. While our development of *OMICmAge* incorporates metabolomics, epigenetics, proteomics, and clinical data, we distilled *OMICmAge* into a single DNAm based aging algorithm through the creation and inclusion of EBP for associated metabolites, proteins, and clinical measures^32–34^. The resulting algorithm, *OMICmAge*, showed several important improvements over previous epigenetic biomarkers. First, *OMICmAge* had stronger associations with all cause mortality relative to the other epigenetic based aging biomarkers we evaluated. We also observed increased accuracy of 10-year death prediction when compared to GrimAge. Notably, we observed strong associations between OMICmAge and major age-related diseases across two distinctly different cohorts. This consistency suggests its broad applicability to population outcomes in a clinical setting. Furthermore, the congruence in our findings remains noteworthy even when considering the use of both ICD9/10 codes and self-diagnoses for prevalent diseases. Besides, high levels of reproducibility has previously been an issue with epigenetic biomarkers which has traditionally only been improved through the inclusion of summary features such as principal components 22^31,32^. With *OMICmAge* alone, we observed high ICCs, demonstrating the strong reproducibility of this metric.

The inclusion of EBPs into the feature space and predictive model for aging biomarkers expands the biological processes that may be represented in epigenetic aging biomarkers. Previously, these biomarkers have had difficulty explaining why aging biomarkers might be accelerated or decelerated in an individual. This has limited the utility of aging biomarkers in clinical practice, as the large heterogeneity in accelerated aging can manifest through many different biological mechanisms. Through the measurement of *OMICmAge* and EBPs, we may better identify the specific biology associated with aging that is captured via the metabolome, proteome, or other clinical factors and assess their overall relationship with *OMICmAge*. The strong correlations between EBPs and the biological measures they are predicting as well as the observed associations with relevant disease and lifestyle factors suggests that some EBPs capture important biology of what they are predicting through DNAm alone.

The relationships between aging and several of the protein and metabolite EBPs selected in *OMICmAge* are well-known. Albumin was selected as the largest weighted EBP in *OMICmAge* and is a protein known to decrease with age^33^. *OMICmAge* also included the androsterone sulfate. Androsterone sulfate is a well-known androgenic steroid that declines with age in men and women due to andropause and menopause that was also retained as a EBP in *OMICmAge*. However, *OMICmAge* also selected proteins and metabolites EBPs with little or no known relationship with aging, such as ribitol which has been identified as a metabolite predictive of mortality but has very little mechanistic information^34^. Furthermore, it is important to recognize that all EBPs and CPGs retained in the predictive model for *OMICmAge* are not causal; nor do they necessarily have the strongest overall associations with *OMICmAge.* They are merely included as predictive variables; further functional work and/or causal modeling via Mendelian Randomization is necessary to infer causality.

There are several limitations which need to be addressed in future work. First is the notion that, while our *EMRAge* model is predicated on EMR data, it’s essential to recognize that real-world data, such as EMRs, might not match the quality of data derived from clinical trials or surveys. This discrepancy arises because EMRs are primarily tailored for clinical care rather than research objectives. The quality of our data was influenced by three primary factors: 1) the absence of complete clinical lab values on the plasma collection date; 2) potential selection bias due to reliance on a singular data source, which could impact the broader applicability of *EMRAge*; and 3) the unavailability of lab test results in the initial study phase, attributed to the limited use of sophisticated analyzers. To mitigate the first concern, we calculated the median lab test value from a five-year span surrounding the collection date, ensuring a robust sample size for *EMRAge* development. The validity of our chosen *EMRAge* predictors was further reinforced by the near-perfect pairwise correlations among four reconstructed *EMRAge* estimates. To address the third limitation, we prioritized commonly conducted lab tests as *EMRAge* predictors. While this strategy might introduce a new selection bias, it effectively maximizes the sample size and maintains a manageable predictor pool. To circumvent the second limitation, we introduced *OMICmAge* and *DNAmAge* models, subsequently corroborating their strong associations with various aging-related diseases across two diverse cohorts. More work will continue to improve the accuracy and precision of EBPs. There are also important considerations with regard to omic data. The profiling platforms for both proteomics and metabolomics do not provide absolute quantification of proteins or metabolites which prevents EBPs from reflecting clinical levels of each variant. Refinement and the use of targeted assays will improve the accuracy of the EBPs. Furthermore, proteomic and metabolomic EBPs could be improved by regressing out known genetic proteinQTL and metaboliteQTL effects from the protein/metabolites levels prior to generating the EBPs. This should be done to preclude the signatures being driven by common SNP data that are invariant across the lifespan. Finally, there is room to expand upon the EBPs that were included into the feature space for *OMICmAge*, both with additional metabolites/proteins and also with other omics. While our omic profiling platforms covered a broad range of metabolites/proteins, we only included 396 EBPs with a significant correlation with *EMRAge*. Future analyses we will expand upon the EBPs in the training of *OMICmAge*. Finally, additional validation of *OMICmAge* across diverse populations will continue to highlight potential limitations in our phenotype. By identifying a healthy biobank cohort, we purposely selected a validation cohort that was distinctly different from MGB-ABC, which had more comorbidities than the general population. We found consistency in the observed associations in our findings, suggesting that *OMICmAge* is robust; however, further interrogation will continue to inform upon our overall understanding of this phenotype.

Overall, we believe the creation of *DNAmEMRAge* and *OMICmAge* represents a step forward in the evolution of improvement of epigenetic aging clocks. *EMRAge* is the first clinical biomarker based clock trained using EMR data to the phenotype of time until death. This provides a unique EMR resource to quantify aging and longevity in large EMR populations. Additionally, *OMICmAge* is the first epigenetic clock that integrates metabolomic, clinical, and proteomic data via EBPs. Finally, *DNAmEMRAge* and *OMICmAge* have the strongest overall associations with prevalent and incidents chronic diseases outcomes and the most accurate 5- and 10-year mortality prediction that were observed.

## Supporting information

Demographics for the study populations used for developing EMRAge, DNAmEMRAge, and OMICmAge and for the study population used for validating the clock

. Information of the 396 Epigenetic Biomarker Proxies (EBPs) included as features in the predictive model for OMICmAge development.

Hazard/Odds ratios of one unit change to disease for multiple aging biomarkers in the MBG-ABC and TruDiagnostic Biobank.

Hazard/Odds ratios of one standard deviation change to disease for multiple aging biomarkers in the MBG-ABC and TruDiagnostic Biobank.

Association between OMICmAge and lifestyle factors in MGB-ABC and TruDiagnostic biobank.

Table of the ICD-9/10 codes.

Table of the extracted clinical phenotypes (N = 27) from the Massachusetts General Brigham (MGB) Biobank.

Table of the selected phenotypes and respective estimated weighting for development of EMRAge.

## Funding/Acknowledgements

Efforts for RK, KM, YC, SB, and JALS are supported by R01HL123915 from the NIH/NHLBI. PK is supported by K99HL159234 from the NIH/NHLBI. Effort for RSK is supported by K01HL146980 from the NIH/NHLBI. Effort for SHC is supported by K01HL153941 from the NIH/NHLBI. Efforts for YC and JAILS are supported by R01HL141826 from the NIH/NHLBI. Effort for AD is supported by K01HL130629 from the NIH/NHLBI. Efforts for AD and JALS are supported by 1R01HL152244 from the NIH/NHLBI. Efforts for MM and JALS are supported by R01HL155742 from the NIH/NHLBI. Effort for MM is supported by R01HL139634 from the NIH/NHLBI. Efforts for JALS, STW, EWK are supported by the NIH U01HG008685. Effort for VNG is supported by NIH AG065403 and AG064223. Efforts for CEW is supported by the Japan Society for the Promotion of Science (JSPS) KAKENHI Grant (JP19K21239), the Japanese Environment Research and Technology Development Fund (No. 5-1752), the Gunma University Initiative for Advanced Research (GIAR), the Japan-Sweden Research Cooperative Program between JSPS and STINT (grant no. JPJ SBP-1201854), the Swedish Heart Lung Foundation (HLF 20180290, HLF 20200693), and the Swedish Research Council (2016-02798). Effort for NJWR is supported by MR/Y010736/1 from the Medical Research Council and IF\R1\231034 from the Royal Society. Effort for EM is supported by the Intramural Research Program of the National Center for Advancing Translational Sciences, NIH (ZICTR000410–03).

## Contributions

QC and VBD had full access to the data and verified the data integrity and accuracy of the analysis.

Conceived and designed the study - JLS, RS

Sample Selection - SC, RK, YC, SB

Performed analysis of physical samples from RPDR - SC, RK, YC, SB, AD, MM, CW, EM

Funding Support of omics - AD, MM, CW, EM, JLS

Performed analysis of physical samples for TruD cohort - TLM, HW

Data processing and normalization - QC, VBD, KM, PK

Algorithm development - QC, VBD, JLS, RS

Statistical analysis and validation - QC, VBD, NCG

Drafted and edited manuscript - all authors

## METHODS

### Discovery Cohort

#### Massachusetts General Brigham (MGB) Biobank

The Massachusetts General Brigham (MGB) Biobank is a large biorepository that provides access to research data and approximately 130,000 high-quality banked samples (plasma, serum, and DNA) from >100,000 consented patients enrolled in the MGB system^35^. These patients can be linked to corresponding Electronic Medical Record (EMR) data, dating from the start of their medical history within the MGB network, in addition to survey data on lifestyle, environment, and family history. The original number of participants from the MGB Biobank who provided plasma samples was 60,371. Of these, 124 participants were excluded because they were younger than 18 years old at the time of plasma collection. Among the remaining adult participants, the vital status of 59,213 has been verified as either alive or deceased, with an accurate record of the date of death as of 07/28/2022. The other 1,034 participants were excluded due to missing verification of their vital status (**Extended Figure S1**).

#### Massachusetts General Brigham Aging Biobank Cohort (MGB-ABC)

The MGB-Aging Biobank Cohort (MGB-ABC) is a cohort of 3,451 randomly selected participants from the MGB Biobank to create a proportionate aging biobank population. The cohort was selected based on even weighting in terms of age, sex and BMI, representative of the MGB Biobank. Comprehensive EMR, metabolomic profiling, proteomic profiling, and epigenetics are available for select subjects in MGB-ABC.

Blood samples, collected either as part of clinical care or through research draws at Brigham and Women’s Hospital (BWH) or Massachusetts General Hospital (MGH), were used for serum, plasma, and DNA/genomic research. Each blood draw typically involved collecting 30-50 ml of blood, which was linked to the corresponding clinical data from the Electronic Medical Record (EMR). The Biobank team also gathered additional health-related information during the blood draw process.

The administration of questionnaires for the study was carried out electronically or in written form, and participants spent approximately 10-15 minutes completing the surveys. The survey included questions related to family history, lifestyle, and environment. and utmost care was taken to ensure the confidentiality and security of the information. Participants’ identities were protected, as no personally identifiable information was requested. The survey data was encrypted to ensure privacy.

The Phenotype Discovery Center (PDC) of MGB integrates various data sources, including the Research Patient Data Registry (RPDR), health information surveys, and genotype results, into the Biobank Portal. This portal combines specimen data with EMR data, creating a comprehensive SQL Server database with a user-friendly web-based application^35^. Researchers can perform queries, visualize longitudinal data with timestamps, employ established algorithms to define phenotypes, utilize automated natural language processing (NLP) tools for analyzing EMR data using the Informatics for Integrating Biology and the Bedside (i2b2) toolkit^36^, and request samples from cases and controls. Data in the Biobank Portal database includes narrative data from clinic notes, text reports (cardiology, pathology, radiology, operative, discharge summaries), codified data (e.g., demographics, diagnoses, procedures, labs and medications) as well as patient-reported data from the health information survey on exposures and family history. Validated phenotypes are available in the Biobank Portal user interface for genotyped Biobank participants. Other relevant measures such as lung function were extracted using a self-developed algorithm incorporating NLP.

#### Metabolomic Profiling

Untargeted global plasma metabolomics profiling was generated by Metabolon Inc. Coefficients of variation were measured in blinded QC samples randomly distributed among study samples. Batch variation was controlled for in the analysis. Sample preparation and global metabolomics profiling was performed according to methods described previously^37^. Metabolomic profiling was performed using four liquid chromatography tandem mass spectrometry (LC-MS) methods that measure complementary sets of metabolite classes described previously^38^: 1) Amines and polar metabolites that ionize in the positive ion mode; 2) Central metabolites and polar metabolites that ionize in the negative ion mode; 3) Polar and non-polar lipids; 4) Free fatty acids, bile acids, and metabolites of intermediate polarity. All reagents and columns for this project will be purchased in bulk from a single lot and all instruments will be calibrated for mass resolution and mass accuracy daily^39^.

Metabolite peaks are quantified using area-under-the-curve. Raw area counts for each metabolite in each sample are normalized to correct for variation resulting from instrument inter-day tuning differences by the median value for each run-day, therefore, setting the medians to 1.0 for each run. Metabolites are identified by automated comparison of the ion features in the experimental samples to a reference library of ~8,000 chemical standard entries that include retention time, molecular weight (m/z), preferred adducts, and in-source fragments as well as associated MS spectra and curated by visual inspection for quality control using software developed at Metabolon, Inc.^39^. Identification of known chemical entities is based on comparison to metabolomic library entries of purified standards. Additional mass spectral entries will be created for structurally unnamed biochemicals, which are identified by virtue of their recurrent nature. These compounds have the potential to be identified by future acquisition of a matching purified standard or by classical structural analysis. Quality control (QC) and data processing was performed using an in-house method that has now been adopted by colleagues across the Boston Longwood Medical Area^40–42^. Briefly, metabolite features with a signal-to-noise ratio <10 were considered unquantifiable and excluded, as were features with undetectable/missing levels for >10% of the samples. All remaining missing values were imputed with the half the minimum peak intensity for that feature across the whole population. Features with a CV in the pooled samples greater than 25% were excluded to ensure good technical reproducibility. Metabolite features were analyzed as measured LC-MS peak areas and were log-transformed to create approximately Gaussian distributions and to stabilize variance, and *pareto* scaled to account for the differences in the scales of measurements across the metabolome. After the QC, 1,459 metabolites across a sample size of 1,986 samples were used in the subsequent analyses.

#### Methylation profiling

DNA methylation/epigenetic data was generated using the Illumina Infinium® MethylationEPIC 850K BeadChip. The MethylationEPIC 850K BeadChip combines comprehensive coverage and high-throughput capabilities with comprehensive genome-wide coverage (greater than 850,000 methylation sites), including CpG islands, non-CpG and differentially methylated sites, FANTOM5 enhancers, ENCODE open chromatin, ENCODE transcription factor binding sites, and miRNA promoter regions. Biobanked samples were stored in −80C prior to shipment to the TruDiagnostic Inc. (Lexington, KY) for DNA extraction and preprocessing. Briefly, 500 ng of DNA was extracted from whole blood samples and bisulfite converted using the Zymo Research EZ DNA methylation kit. All manufacturer’s instructions were followed. Bisulfite-converted DNA was randomly assigned to chip wells on the Infinium HumanMethylationEPIC array. Lab preprocessing included the following: 1) DNA amplification, 2) hybridization to the EPIC array, 3) stain, washing, and imaging with the Illumina iScan SQ instrument to generate raw image intensities.

Raw methylation data for the MGB-Biobank was processed using the *minfi* pipeline^43^, and low quality samples were identified using the qcfilter() function from the *ENmix* package^44^, using default parameters. Overall, a total of 4,803 samples passed the QA/QC (p < 0.05) and were deemed to be high quality samples. In addition, we removed low quality probes (p < 0.05 out-of-band) that were identified among the samples. This process retained 721,802 among 866,239 probes that were high quality and indicated that the large portion of the methylation data was of high quality. A combinatorial normalization processing using the Funnorm procedure (*minfi* package), followed by RCP method (*ENmix* package) was performed in order to minimize sample to sample variation as noted in Foox et al. 2021^45^.

#### Proteomic Profiling

We used the Seer proteomic platform for its ability to discover novel proteins and peptides related to chronological and biological aging. While Seer uses LC-MS/MS like other proteomic platforms, its patented and novel nanoparticles strategy uses nanoparticles with different binding capabilities to isolate and extract peptides and proteins via corona covalent attachment to its surface. This technique is unique as it gives the platform the ability to detect low abundance peptides and proteins without the need to depletion of the fraction which can be 10x more expensive. Moreover, this increases the number of quantified proteins by 3–5 fold compared to depleted and un-depleted serum proteomics. The Proteograph untargeted approach is also different from other popular platforms based on aptamer technologies which only quantifies peptides which they specifically target. This also allows for an unbiased discovery of proteomic associations.

Relative protein levels were quantified for 2,000 samples - 1,600 from the MGB-ABC and 400 process controls - using the Proteograph Product Suite (Seer, Inc.) and LC-MS. Briefly, the samples were incubated with five proprietary nanoparticles which formed protein coronas on the Seer SP100 proteograph, which allowed for the capture of proteins using physicochemical binding. The resulting proteins were digested using trypsin, and relative levels were quantified using the default DIA method provided in the Protograph Analysis Software (PAS). Protein groups were ultimately considered for downstream analyses for two reasons: peptides identified can ambiguously map to multiple proteins, and combining peptides into protein groups can improve protein quantification devoid of spurious quantification. Thus, protein group data was sent through pre-processing, control based normalization, and outlier detection. This produced estimations of a total of 28,490 peptides across blood samples (average 15,239), and 10,265 (average 4,281) across the controls. Peptides were then further consolidated into 3,695 total protein groups (average 2,587) in MGB-ABC samples, and 1,360 total protein groups among the plate controls. Following the signal drift and batch effect correction via the Quality Control-Robust Spline Correction (QCRSC) algorithm^46^, we applied Log10 Transformation, Pareto Scaling, and kNN imputation based on current guidelines^47^. Stringent filters, including 80% protein presence, RSD-qc < 0.20%, and D-ratio < 0.70, were utilized to reinforce data validity and reliability^48^. The final processed dataset consists of 2,805 non-unique, or 536 unique, protein groups, across 1,789 samples, in which the majority of samples (N = 1,475) matched to methylation data.

#### Definition of age-related diseases

We utilized ICD-9/10 codes to identify age-related diseases, including type-2 diabetes, COPD, depression, cancer, stroke, and other cardiovascular diseases, as detailed in **Extended Table S5**.

### Validation cohort

#### TruDiagnostic Biobank cohort

The TruDiagnostic Biobank cohort included 13,109 individuals who took the commercial TruDiagnostic TruAge test and had their DNA methylation data generated. The participants were recruited between October 2020 and April 2023 and were predominantly from the United States. These participants were in better health compared with individuals from Mass General Biobank, likely due to their proactive interest in health and willingness to pay for epigenetic testing. The majority of these samples were performed under a healthcare provider’s recommendation and guidance while less than 5% were in a direct-to-consumer setting. As a result, these individuals may experience a self selection bias whereby they seek preventative medicine and have fewer comorbidities than normal patient populations. During the recruitment of participants, they were asked to complete a survey that included questions about personal information, medical history, social history, lifestyle, and family history. The study involving human participants was reviewed and approved by the IRCM IRB and the participants provided written informed consent to take part in the study.

#### Methylation profiling

Peripheral blood samples were collected using a lancet and capillary method and placed in a lysis buffer for DNA extraction. Then, 500 ng of DNA was treated with bisulfite using the EZ DNA Methylation kit from Zymo Research following the manufacturer’s instructions. The bisulfite-treated DNA samples were randomly assigned to a well on the Infinium HumanMethylationEPIC BeadChip, which was then amplified, hybridized, stained, washed, and imaged with the Illumina iScan SQ instrument to obtain raw image intensities.

To pre-process the TruDiagnostic methylation data, we used the same pipeline as for the MGB-ABC cohort. A total of 12,666 individuals, representing 96.7% of the original samples, passed the QA/QC (p < 0.05) and were deemed to be high quality samples. However, we did not remove any probe in order to keep all the CpG sites needed for clock calculation. Due to computational limitations, we were unable to apply the same normalization methods as in the MGB-ABC cohort. Thus, we applied normal-exponential out-of-band (Noob) normalization using the *preprocessNoob* function from the *minfi* package. Finally, we used a 12 cell immune deconvolution method to estimate cell type proportions^49,50^.

### Statistical analysis

#### Development of EMRAge

We extracted 27 clinical phenotypes from 59,213 participants in the MGB Biobank with plasma samples (**Extended Table S6**). To address missing values and instrumental variations, the median value of all numerical observations, except height and age, were replaced with missing median values of all median observations within 5 years around the first plasma collection. The resulting individuals with complete data for all clinical phenotypes were used in the analysis (n=28,733). Selected samples were divided into training and testing sets using a 70:30 ratio, and then fitted a Cox proportional hazards (Cox-PH) model in the training set to estimate the weightings (i.e., coefficients) of the selected features. To assure the generalizability of the equation, we did not scale the training data. Using the trained model, we calculated the risk estimate for each individual in the training set by linearly combining the estimated weightings and predictor values (*Xβ*). This estimate was further transformed into the *EMRAge* value using the equation below to get the same mean and variance as chronological age:

**EMRAge = 9.95006 * Xβ_{train} + 52.14512**

Where 9.95006 belongs to the mean of chronological age and 52.14512 to the standard deviation.

We utilized one-hot encoding to convert categorical variables into binary variables and standardized the numerical variables. Then we used a LASSO Cox regression model to predict time-til-death among 28,733 individuals. To determine the optimal model, we assessed model performance based on Harrell’s C index. As a result, the optimized model selected 25 clinical variables, which were then passed to a Cox Proportional-Hazard (Cox-PH) model for further filtering. Ultimately, 19 clinical variables remained as they had an adjusted p-value ≤ 0.05 in the fitted Cox-PH model. As a result, 30,884 participants with all available electronic medical records of selected clinical variables remained in the final cohort. We then evaluated the Pearson correlation of *EMRAge* to chronological age in the training and testing sets.

Since the *EMRAge* predictors (i.e., clinical phenotypes) were selected based on imputed data, we validated their robustness by re-training the algorithm at four different time points: Jan 1st of 2008, 2010, 2012, and 2014. During each time point, we replaced the values of the selected phenotypes with the median values of observations within a one-year timeframe, ensuring no overlap of data scans. To make a sensible comparison, we excluded the Charlson Comorbidity Index indicators because they are less sensitive to time in our data. As a result, we had 4 estimating equations trained using the same predictors but different data. Then we applied these equations to calculate the *EMRAge* value of the participants (N=11,673) with complete data of predictors around 1-year centered on January 1st of 2016, and then checked Pearson’s correlations among these four estimates.

We then assessed the relationship between *EMRAge* and both incident and prevalent aging-related health outcomes, including all-cause mortality, stroke, type-2 diabetes, COPD, depression, other cardiovascular diseases (CVD), and any type of cancer. We used the following criteria based from the patient EMRs for a positive diagnosis of an age-related health outcome: 1) at least 2 relevant ICD-9/10 codes were recorded in the patient’s EMR, and 2) the first and last dates of the relevant ICD codes should be at least 1 day apart. We estimated the incident risk of the *EMRAge* for all prospective adverse events in the Cox-PH model, adjusting for age, sex, race, BMI, smoking status and alcohol consumption. We also estimated the odds ratio of the *EMRAge* for all prevalent morbidities in the logistic regression model, adjusting for the same covariates.

#### Comparison of EMRAge and PhenoAge

We compared the incidence and prevalence of age-related health outcomes between *EMRAge* and PhenoAge by applying the same linear models described above with PhenoAge as the outcome. We calculated PhenoAge for our MGB Biobank cohort using the established toolkit, as initially proposed by Levine et al. (2018) and developed by Belsky and Kwon^51^. Although PhenoAge is derived from eight clinical lab metrics, one specific parameter, C-reactive protein, is not frequently ordered in routine clinical settings. To maximize sample retention, we employed the same strategy to retain the median value of observations over a 5-year window centered on the plasma collection date. Following this imputation, our sub-cohort consisted of 17,093 participants, with 11,945 samples in the training set and 5,148 in the testing set.

#### Development of DNAmEMRAge

After developing the *EMRAge* measure, we next created a DNAm surrogate predictor of *EMRAge* using matched EPIC array data (*DNAmEMRAge*). To this end, we used the MGB-ABC cohort, which is a subset of the MGB Biobank that was created with the aim of possessing a proportionate aging biobank population. DNA Methylation was generated from a total of 4,803 samples using the EPICv1 array. To allow for training, samples were then selected for having *EMRAge* quantified and the availability of chronological age and sex information, which retained 3,451 samples. To develop *DNAmEMRAge*, the normalized DNA methylation dataset was transposed so that CpG sites were considered features, and trained to *EMRAge* values. Using an 80-20 train to test split, the *glmnet* R package^52^ (version 4.1-8) was used to train a Gaussian penalized regression model using an alpha parameter of 0.1. Using a cross validation fold number of 25, an optimal lambda was selected based on the alpha parameter. Sex was classified as Gender_M (males) and Gender_F (females) using one-hot encoding, and was included as penalized features along with chronological age and the relative cell proportions of 12 immune cell types. All CpGs and the covariates mentioned were included as penalized features. Those features that showed a non zero coefficient were selected for the final model.

#### Development of Epigenetic Biomarker Proxies (EBP) models

To generate EBPs elastic net models, the normalized DNA methylation dataset was transposed to which the features were considered to be CpG sites. CpGs were not pre-filtered, and all CpGs that passed QC were used for training. Training was conducted to each significant clinical, metabolite, and protein group values. To ensure the best model associated with *EMRAge* was generated, the *glmnet* R package was used to train Gaussian penalized regression models across all measures. In order to maximize the number of CpGs, an alpha value of 0.1 was utilized. However, for three clinical values (smoking pack years, total bilirubin, and total cholesterol), an alpha of 0.5 was applied as they produced the highest correlation between predicted and observed values. Using a cross validation fold number of 25, an optimal lambda was selected at each alpha threshold and implemented to select the features. Sex was classified as Gender_M (males) and Gender_F (females) using one-hot encoding, and was included as penalized features along with chronological age. Features that showed a non zero coefficient were selected for each EBP. EBPs with a significant (p<0.05) Pearson correlation greater than 0.2 were selected to include as features in the model to train *OMICmAge*.

We assessed the association between the EBPs and multiple chronic disease and lifestyle outcomes in both MGB-ABC and TruDiagnostic Biobank cohort. Specifically, we standardized all EBPs and utilized a generalized linear regression model with disease/lifestyle as the outcome and each individual EBP as the predictor variable, adjusting for age, sex, ethnicity, BMI, and tobacco smoking.

#### Development of OMICmAge

*OMICmAge* was developed using the individuals used to develop *DNAmEMRAge* (N = 3,451), and applying a similar 80:20 train:test split. Penalized regression models using an alpha of 0.1 and the optimal lambda identified after a 25 cross-validation were used to train a single composite model among the train samples. For training, the following values were inputted as penalized features regression model and trained to *EMRAge*: all CpG sites present in the first and second generation of Illumina EPIC arrays; all selected clinical, metabolite, and proteins EBPs estimates from the training samples; relative estimates of 12 cell immune cell subtypes; and the demographic information (Age, sex, and BMI). For sex, a similar one-hot encoding was used to identify males (Gender_M) and females (Gender_F). Features which showed non-0 coefficients were kept in the final multivariate model.

#### Comparison of OMICmAge, EMRAge and Previous Clocks

For comparison of *OMICmAge* and *EMRage* clocks to previous methods of biological age prediction, we chose to analyze PCHorvath^10^, PCHannum^11^, PCPhenoAge^51^, PCGrimAge^20^, and DunedinPACE^53^. We chose their PC (i.e., principal component) versions as they have much better precision while still maintaining their relationships to health outcomes^54^. In order to compare the CpG sites included in each model, we used the non-PC clocks because the PC models do not contain CpG sites as predictors.

We used the Cox Proportional-Hazard regression model to assess the association between each clock and incident age-related diseases, such as type-2 diabetes, stroke, depression, COPD, other cardiovascular diseases, and any type of cancer. We also employed the logistic regression model to assess the association between each clock and prevalent age-related diseases. Each model was adjusted for age, gender, race, BMI, smoking status, and alcohol drinking habits. Furthermore, to predict the 5-year and 10-year survival probability, we used a simple logistic regression model with a binary survival flag as the outcome and each clock as the sole predictor. We drew ROC curves and estimated the Area Under the Curve (AUC) based on this model.

---

**Extended Figure S1.**
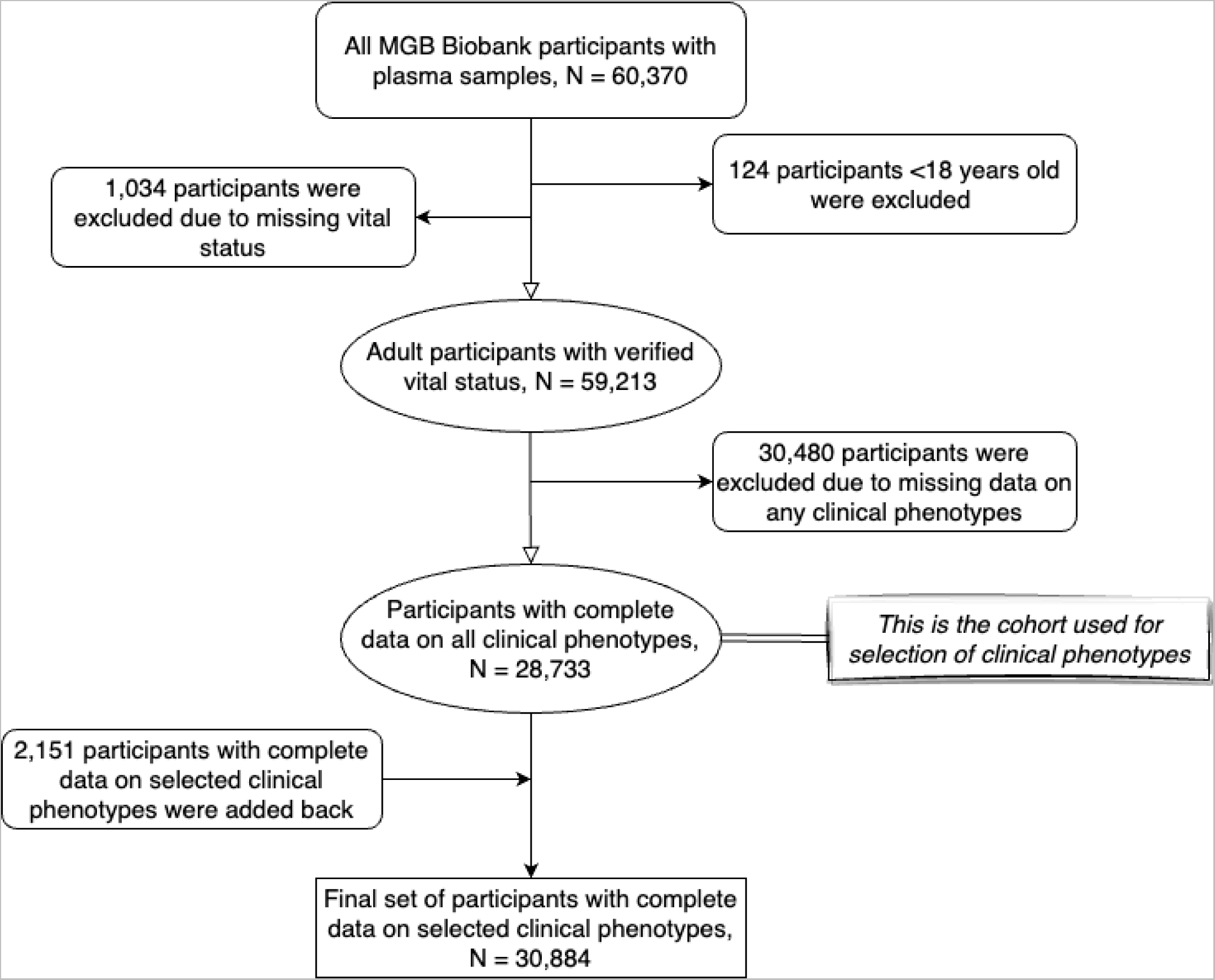
Flowchart for inclusion of participants from the Massachusetts General Brigham (MGB) Biobank for the development of *EMRAge*.

**Extended Figure S2.**
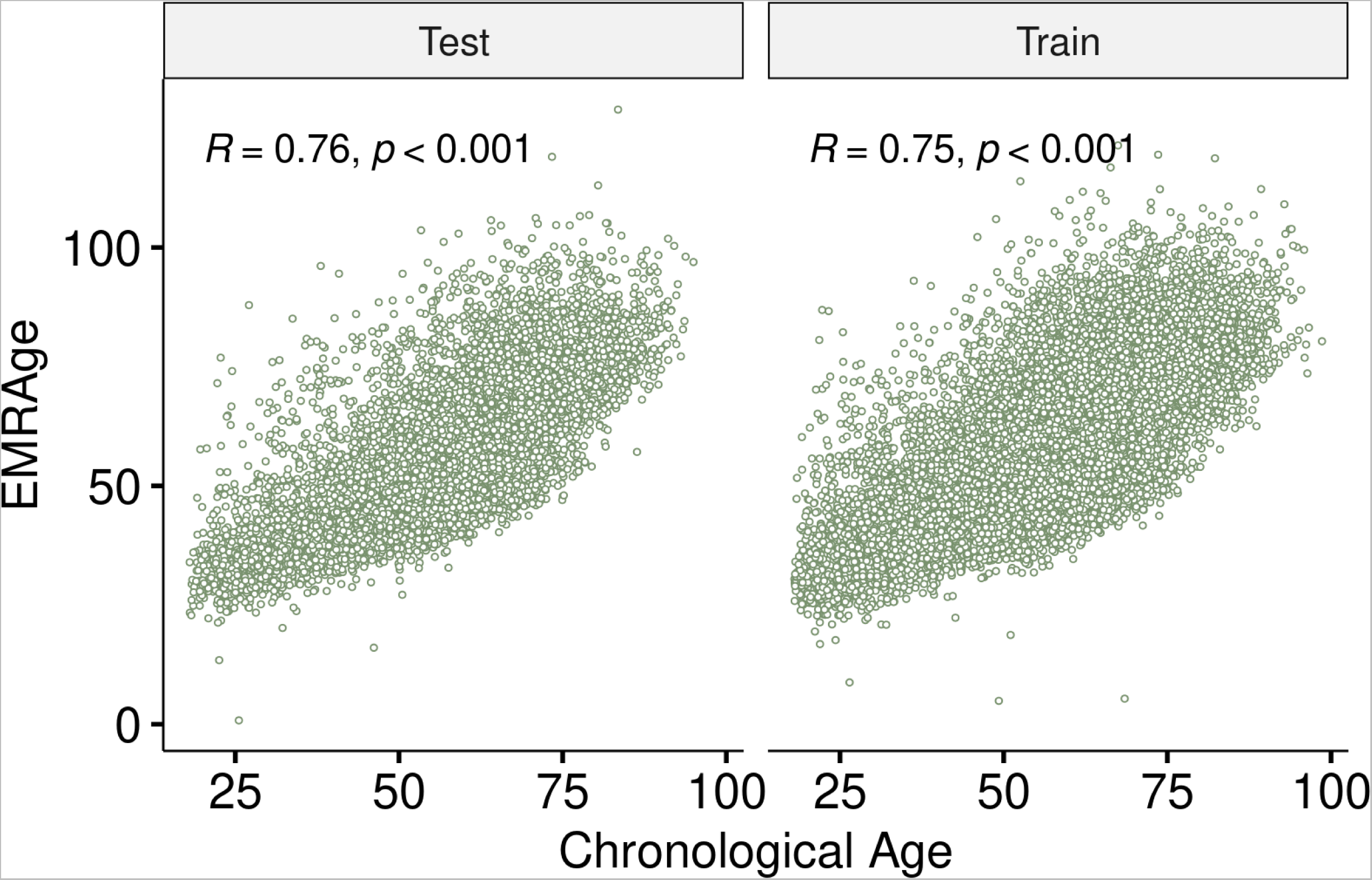
Pearson’s correlation between *EMRAge* and chronological age in the test and train sets.

**Extended Figure S3.**
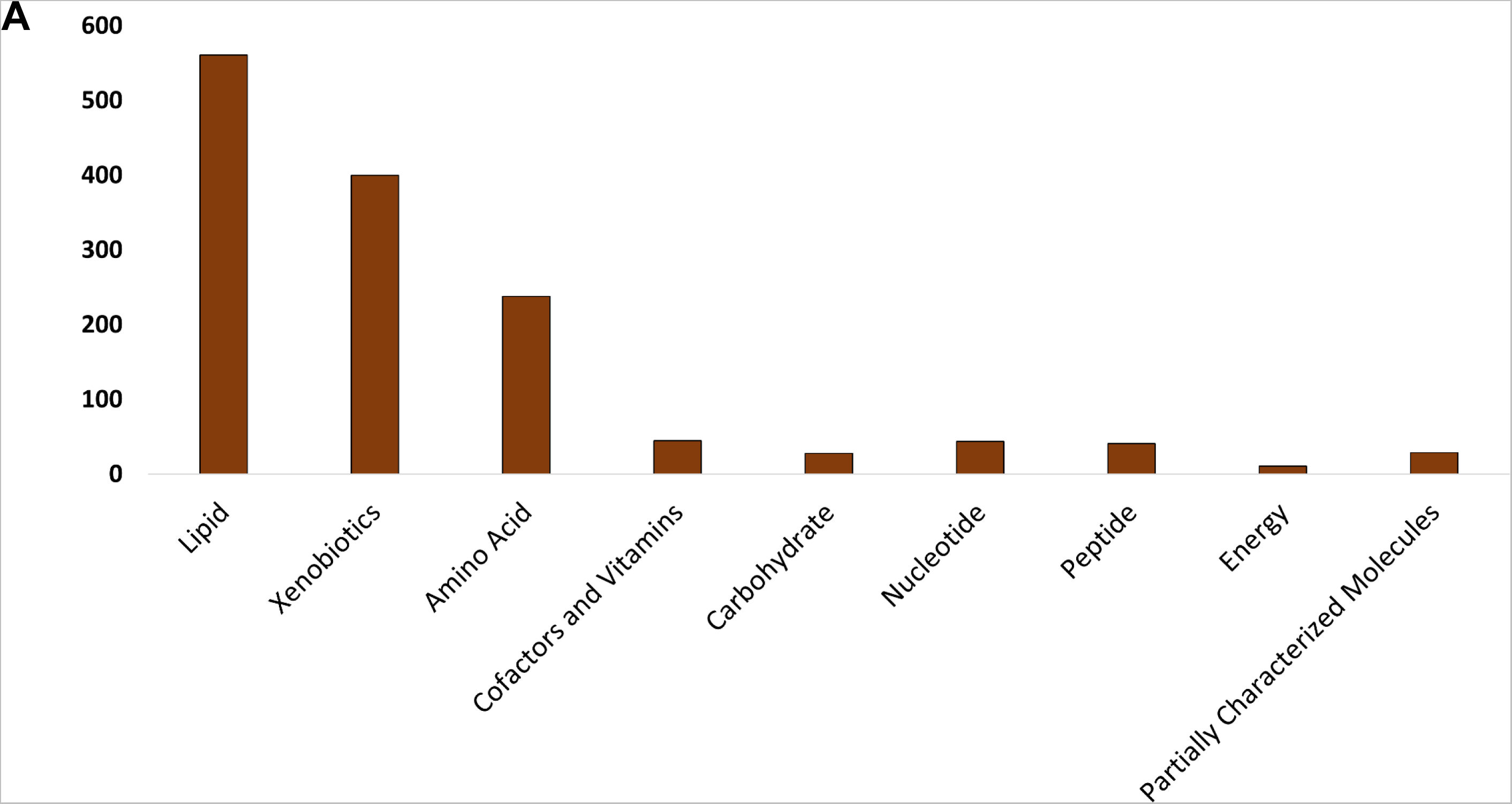
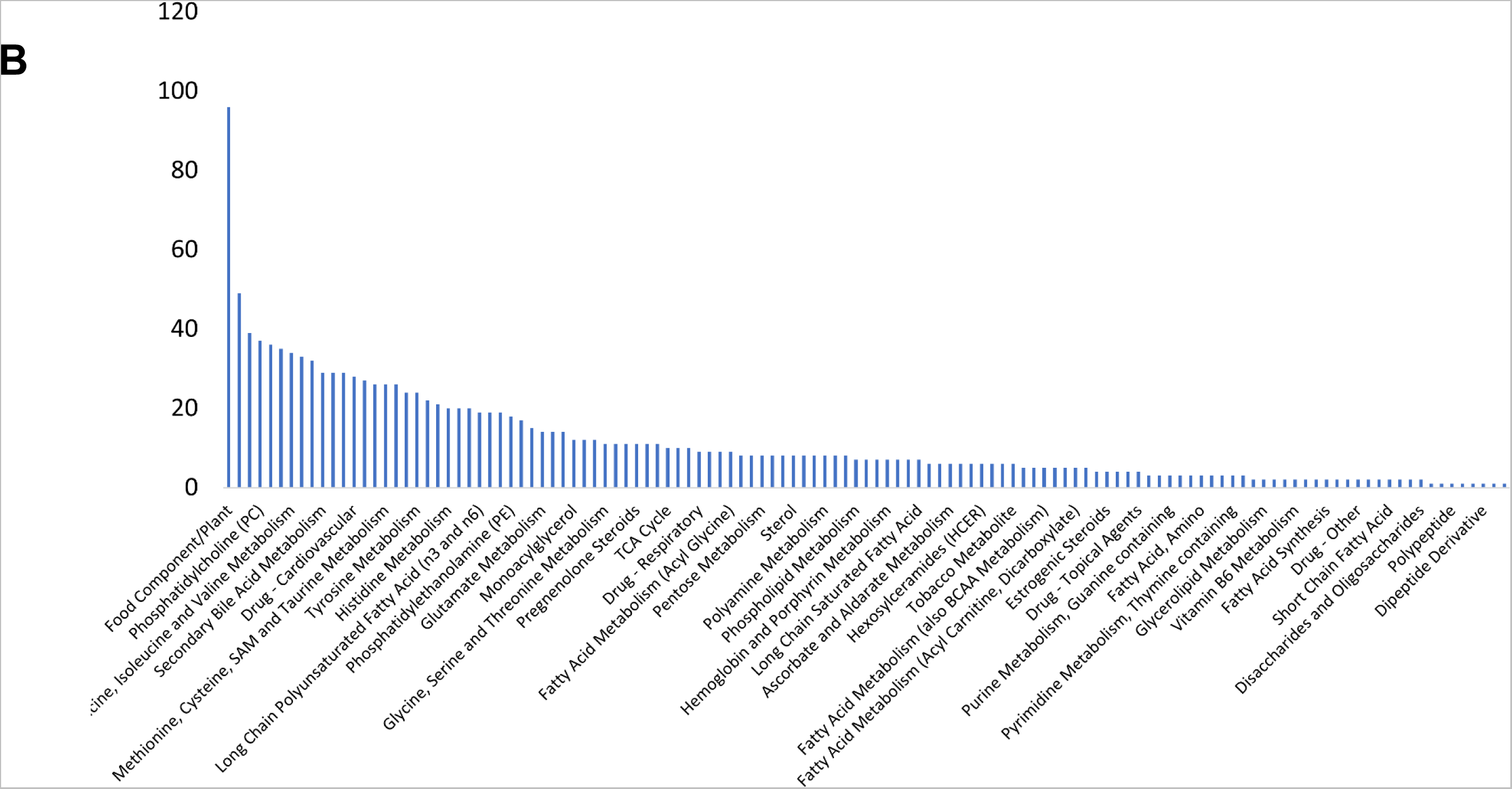
Distribution of super pathways (A) and subpathways for all the metabolites (B).

**Extended Figure S4.**
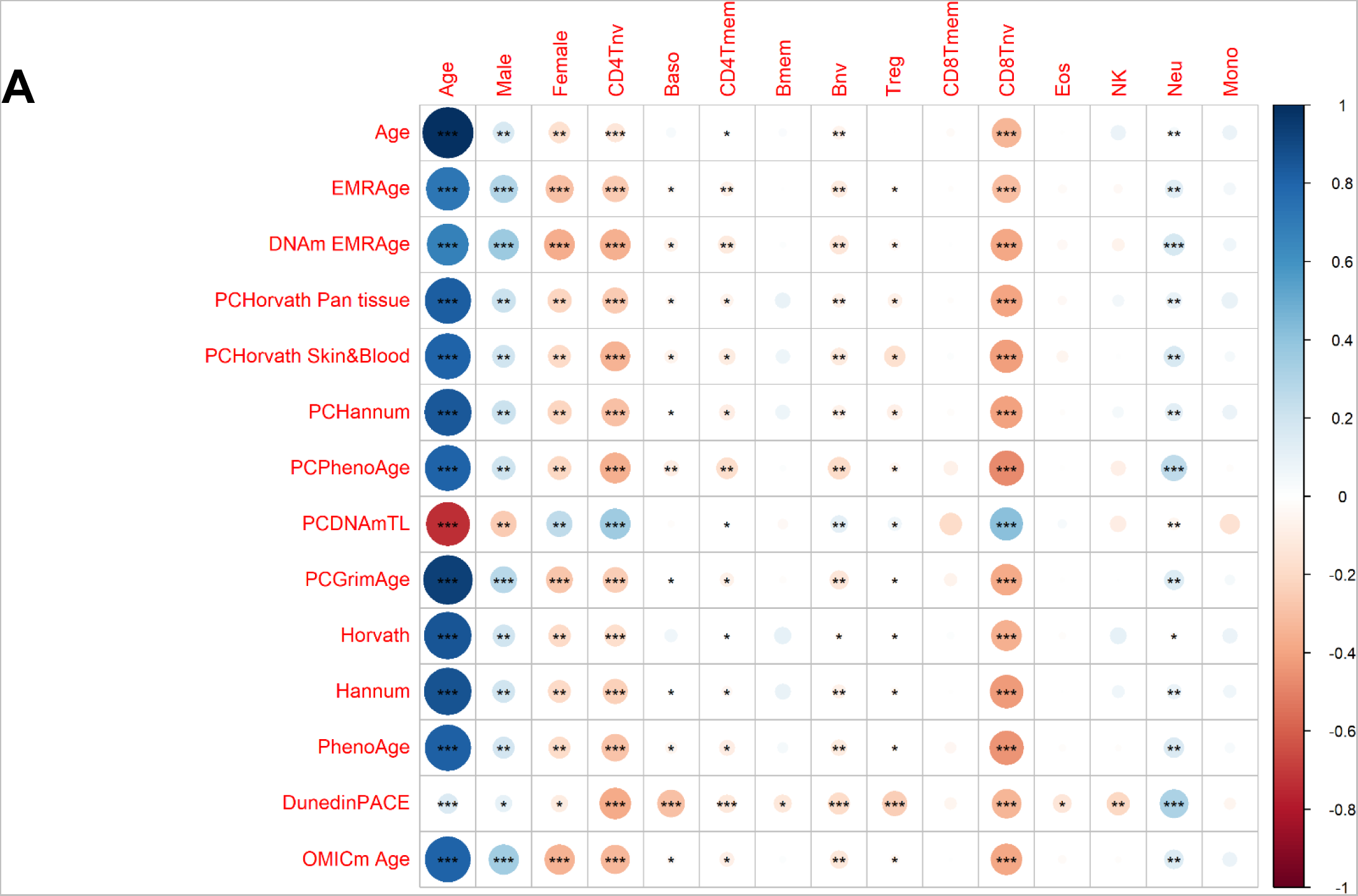
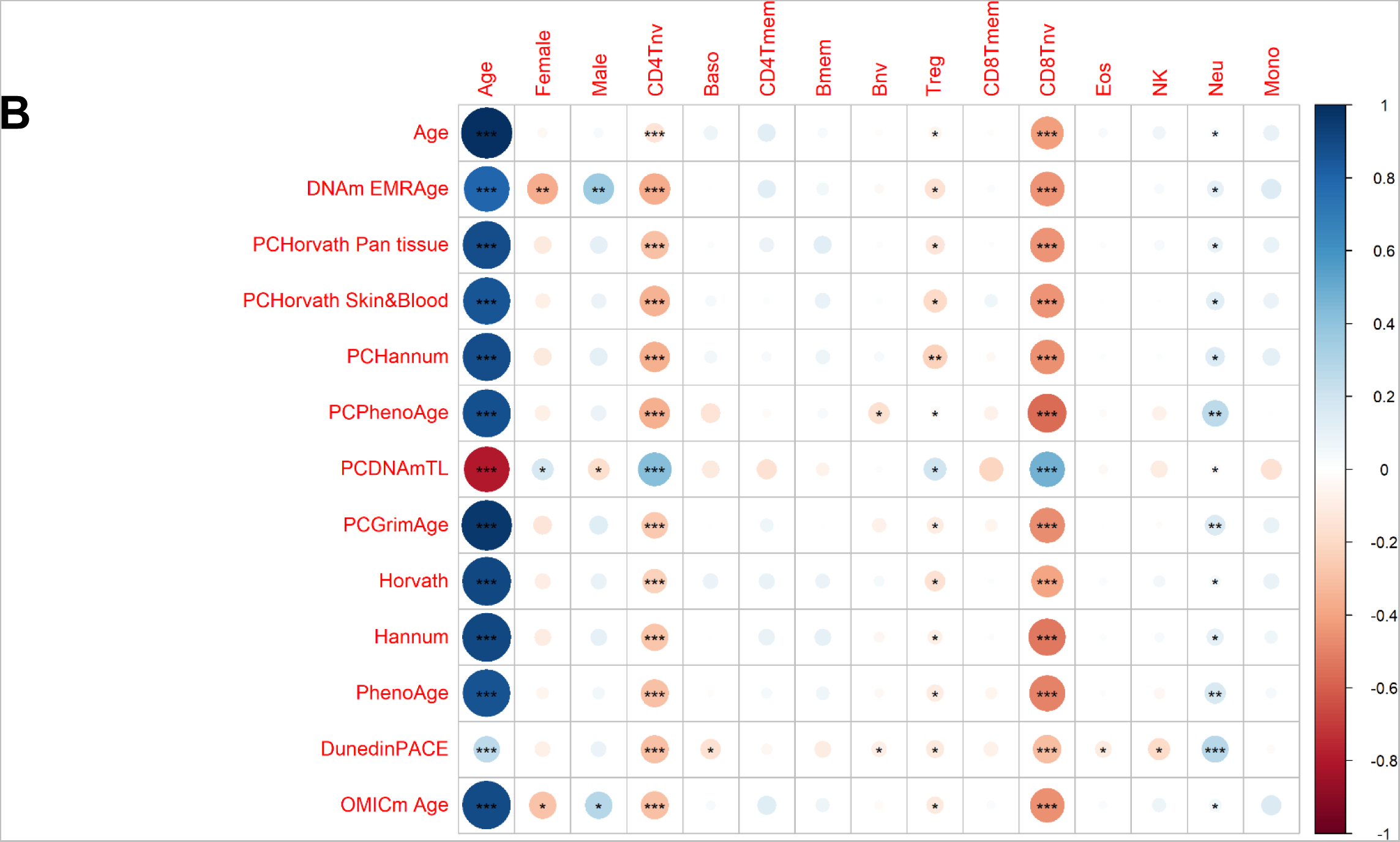
Correlation plots between epigenetic clocks and immune cells in the MGB-ABC and TruDiagnostic biobanks. A) Discovery cohort - MGB-ABC biobank. B) Validation cohort - TruDiagnostic biobank. Each column represents a covariate among age, gender male, gender female, and immune cells. The size is related to the magnitude of the correlation and the color to the direction (positive or negative). *: p-value < 0.05, **: p-value < 0.01, ***: p-value < 0.001.

## Extended Tables

**Extended Table S1. Demographics for the study populations used for developing *EMRAge*, *DNAmEMRAge*, and *OMICmAge* and for the study population used for validating the clocks.**

**Extended Table S2. Information of the 396 Epigenetic Biomarker Proxies (EBPs) included as features in the predictive model for *OMICmAge* development.** Among them, 266 are metabolite EBPs, 109 are protein EBPs, and 21 are clinical EBPs. The table contains information on MSE, the R^2^, and the Pearson correlation between the EBPs and the values for each feature. For those EBPs selected for the *OMICmAge* after penalization, there is also a description of the biomarker.

**Extended Table S3a. Hazard/Odds ratios of one unit change to disease for multiple aging biomarkers in the MBG-ABC and TruDiagnostic Biobank.**

**Extended Table S3b. Hazard/Odds ratios of one standard deviation change to disease for multiple aging biomarkers in the MBG-ABC and TruDiagnostic Biobank.**

**Extended Table S4. Association between *OMICmAge* and lifestyle factors in MGB-ABC and TruDiagnostic biobank3**

**Extended Table S5. Table of the ICD-9/10 codes**

**Extended Table S6. Table of the extracted clinical phenotypes (N = 27) from the Massachusetts General Brigham (MGB) Biobank.**

**Extended Table S7. Table of the selected phenotypes and respective estimated weighting for development of EMRAge**

