## Supplementary material for "OMICmAge: An integrative multi-omics approach to quantify biological age with electronic medical records": Demographics for the study populations used for developing EMRAge, DNAmEMRAge, and OMICmAge and for the study population used for validating the clock

**Extended Table S1. Demographics for the study populations used for developing *EMRAge*, *DNAmEMRAge*, and *OMICmAge* and for the study population used for validating the clocks.**

|  | ***EMRAge*** | ***DNAmEMRAge***  ***OMICmAge*** | **Biomarkers validation** |
| --- | --- | --- | --- |
| **Study population** | **MGB Biobank** | **MGB - Aging Biobank Cohort** | **TruDiagnostic Biobank** |
| **N** | 30,884 | 3,451 | 12,666 |
| **Sex, male, N (%)** | 13,759 (44.6%) | 1,250 (36.2%) | 7486 (59.1%) |
| **Age in years, mean (range)** | 56.6 (18.0 - 98.6) | 58.5 (18.3 - 96.3) | 53.2 (12.9-100) |
| **Race/Ethnicity, N (%)** |  |  |  |
| White | 25,940 (84.0%) | 2,802 (81.2%) | 9718 (76.7%) |
| African American | 1,785 (5.8%) | 250 (7.2%) | 184 (1.5%) |
| Asian | 710 (2.3%) | 59 (1.7%) | 744 (5.9%) |
| Other | 1,700 (5.5%) | 263 (7.6%) | 2020 (15.9%) |
| Unknown/Missing | 749 (2.4%) | 77 (2.2%) | 0 (0%) |
| **BMI* (kg/m^2^), mean (range)** | 28.7 (12.6 - 83.0) | 30.0 (15.5, 69.0) | 25.4 (6.9-60.3) |
| **Tobacco Use, N (%)** |  |  |  |
| Yes | 1,752 (5.7%) | 212 (6.1%) | 537 (4.2%) |
| No* | 29,132 (94.3%) | 3,239 (93.9%) | 11918 (94.1%) |
| Missing | 0 (0%) | 0 (0%) | 214 (1.7%) |
| **Alcohol Drink, N (%)** |  |  |  |
| Yes | 17,941 (58.1%) | 1,619 (46.9%) | 10,086 (79.6%) |
| No | 11,516 (37.3%) | 1,761 (51.0%) | 2,366 (18.7%) |
| Missing | 1,427 (4.6%) | 71 (2.1%) | 214 (1.7%) |
| **Prevalence of comorbidity at baseline, N (%)** |  |  |  |
| Stroke | 2,073 (6.7%) | 286 (8.3%) | 100 (0.8%) |
| Type-2 Diabetes | 5,582 (18.1%) | 1,139 (33.0%) | 258 (2.0%) |
| COPD | 2,391 (7.7%) | 865 (25.1%) | 59 (0.5%) |
| Depression | 8,572 (27.8%) | 1,570 (45.5%) | 44 (0.3%) |
| Cancer | 5,539 (17.9%) | 691 (20.0%) | 1301 (10.3%) |
| CVD (excl. stroke) | 10,501 (34.0%) | 1,892 (54.8%) | 4301 (33.9%) |
| **Vital Status, as of Jul 28 2023, N (%)** |  |  |  |
| Deceased | 2,405 (7.8%) | 418 (12.1%) | - |
| Alive | 28,479 (92.2%) | 2,886 (83.6%) | - |
| Unknown | - | 147 (4.3%) | - |

*BMI: Body Mass Index. BMI is based on median values of BMI records within 5 years around the first plasma collection for the MGB Biobank samples.

*For tobacco use, the “No” level combines the non-smokers and the former smokers.
