## Supplementary material for "OMICmAge: An integrative multi-omics approach to quantify biological age with electronic medical records": Table of the extracted clinical phenotypes (N = 27) from the Massachusetts General Brigham (MGB) Biobank.

**Extended Table S6. Table of the extracted clinical phenotypes (N = 27) from the Massachusetts General Brigham (MGB) Biobank.**

Demographics

Gender (Female / Male)

Race (White / Black / Asian / Other)

Age at plasma collection

Risk Index

Charlson Comorbidity Index (0 / 1 / 2 / 3 / 4 / 5 / 6 / 7+)

Substance Use

Smoking status (Non-smoker / Former-smoker / Current-smoker)

Alcohol drinking (No-alcohol / Former-drinker / Current-drinker)

Vital Sign

Height (cm)

BMI (kg/m^2)

Systolic Blood Pressure (mmHg)

Diastolic Blood Pressure (mmHg)

Basic Metabolic Panel

Blood Urea Nitrogen (mg/dL)

Creatinine (mg/dL)

Glucose (mg/dL)

Hepatic Function Panel

Albumin (g/dL)

Total Bilirubin (mg/dL)

Alkaline phosphatase (U/L)

Alanine transaminase (U/L)

Aspartate transaminase (U/L)

Lipid Panel

High-density Lipoprotein (mg/dL)

Low-density Lipoprotein (mg/dL)

Triglyceride (mg/dL)

CBC and Differential

Hematocrit (%)

Platelet Count (10^9/L)

Red Cell Width Distribution

White Blood Cell count (10^9/L)

Basophils count (K/uL)

Eosinophils count (K/uL)

Neutrophils count (K/uL)
