## Supplementary material for "OMICmAge: An integrative multi-omics approach to quantify biological age with electronic medical records": Table of the selected phenotypes and respective estimated weighting for development of EMRAge.

**Extended Table S7. Table of the selected phenotypes and respective estimated weighting for development of EMRAge**

| **Clinical Variable** | **Estimated Weighting** |
| --- | --- |
| Age | 0.0374 |
| BMI | -0.0148 |
| Blood Urea Nitrogen | 0.0113 |
| Charlson Comorbidity Index - 6 | 0.8261 |
| Charlson Comorbidity Index - 7+ | 1.2362 |
| Diastolic Blood Pressure | 1.1341 |
| Eosinophil count | -0.0178 |
| Female | -0.0064 |
| Glucose | -0.3938 |
| Hematocrit | 0.0047 |
| Albumin | -0.0405 |
| Alkaline phosphatase | -0.7805 |
| Alanine transaminase | 0.0021 |
| Aspartate transaminase | -0.0073 |
| Neutrophil count | 0.0051 |
| Platelet count | 0.0004 |
| Red Cell Distribution Width | -0.0015 |
| Current-smoker | 0.2021 |
| Triglyceride | 0.2355 |
